# Structural basis of undecaprenyl phosphate glycosylation leading to polymyxin resistance in Gram-negative bacteria

**DOI:** 10.1101/2025.01.29.634835

**Authors:** Khuram U Ashraf, Mariana Bunoro-Batista, T. Bertie Ansell, Ankita Punetha, Stephannie Rosario-Garrido, Emre Firlar, Jason T. Kaelber, Phillip J. Stansfeld, Vasileios I. Petrou

## Abstract

In Gram-negative bacteria, the enzymatic modification of Lipid A with aminoarabinose (L-Ara4N) leads to resistance against polymyxin antibiotics and cationic antimicrobial peptides. ArnC, an integral membrane glycosyltransferase, attaches a formylated form of aminoarabinose to the lipid undecaprenyl phosphate, enabling its association with the bacterial inner membrane. Here, we present cryo-electron microscopy structures of ArnC from *S. enterica* in *apo* and nucleotide-bound conformations. These structures reveal a conformational transition that takes place upon binding of the partial donor substrate. Using coarse-grained and atomistic simulations, we provide insights into substrate coordination before and during catalysis, and we propose a catalytic mechanism that may operate on all similar metal-dependent polyprenyl phosphate glycosyltransferases. The reported structures provide a new target for drug design aiming to combat polymyxin resistance.

## Introduction

Antibiotic resistance is a growing global public health threat that endangers our ability to treat common infections^1^. In 2019, the Centers for Disease Control and Prevention (CDC) estimated that more than 2.8 million antibiotic resistant infections occur in the United States every year, leading to more than 35,000 deaths^2^. The situation is particularly critical with the emergence of multidrug resistant (MDR) Gram-negative (GN) strains of pathogens, including *Klebsiella pneumoniae*, *Acinetobacter baumanii*, *Pseudomonas aeruginosa*, *Salmonella enterica*, and *E. coli*, which are responsible for MDR nosocomial and non-nosocomial infections^3,4^.

Polymyxins are currently the last line of defense against MDR Gram-negative bacterial infections^5–7^. Polymyxins are non-ribosomal lipopeptides that are polycationic at physiological pH^8^. They act by permeabilizing bacterial membranes^9^, although alternative mechanisms like lipid exchange have been proposed^10,11^. In all cases, the necessary initial step is the binding of polymyxins to the lipopolysaccharide (LPS) of the outer membrane^12^ (**Fig. 1A**). This molecular association is achieved through electrostatic interactions between the cationic amino groups of polymyxins and anionic phosphate groups of Lipid A, which is the amphipathic saccharolipid that anchors LPS to the membrane^12^. Resistance to polymyxins can emerge spontaneously *in vitro*^13–15^ and has been observed in patients in case of suboptimal use^16,17^. The resistance is mainly acquired through enzymatic modifications of Lipid A that cap the glucosamine sugar phosphates, resulting in a reduction of the negative charge in bacterial outer membranes^18^. This mechanism also represents an evasion tactic of GN bacteria against natural antimicrobial peptides (AMPs), including those generated by the innate immune system^19,20^. Clinical and non-clinical isolates that exhibit resistance to polymyxins often carry chromosomal mutations in control elements that allow upregulation of Lipid A-modifying enzymes^21^.

**Figure 1.**
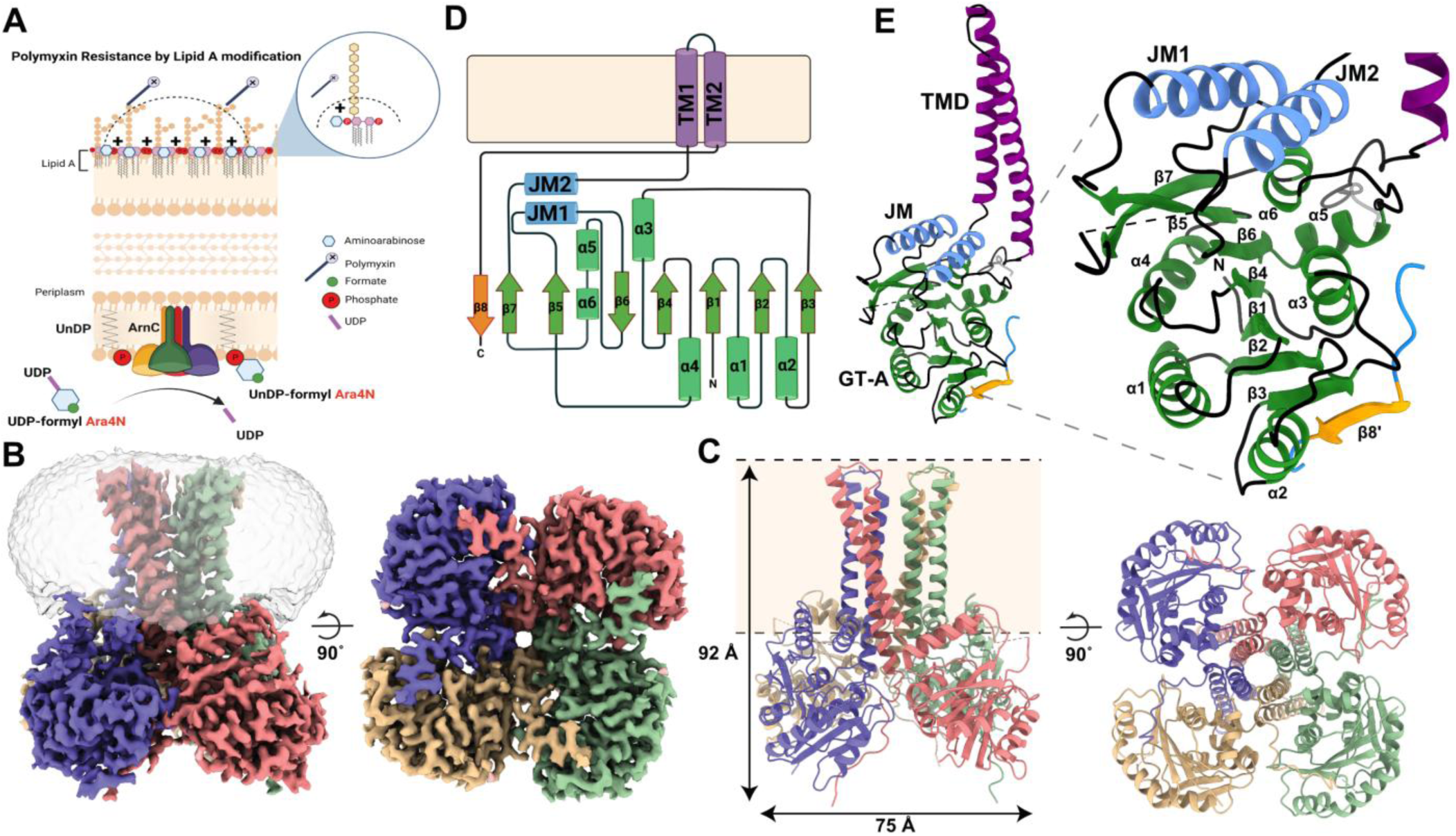
The cryo-EM structure of ArnC from *S. enterica*. **A**) Schematic representation of the ArnC enzyme in the inner membrane of Gram-negative bacteria and the chemical reaction it catalyzes. The aminoarabinose sugar after its association with UndP is ultimately used to decorate the phosphates of Lipid A, reducing the negative charge of the outer bacterial membrane, and leading to polymyxin resistance. **B**) Cryo-EM density map of the ArnC tetramer from the Krios dataset in two orthogonal views. Coloring is per protomer, and the semi-transparent volume represents the nanodisc density observed in low contour levels of the reconstruction. **C**) The cryo-EM structure of ArnC in ribbon representation. Approximate membrane boundaries are represented with dotted lines. The coloring is per protomer, and two orthogonal views are shown (side and bottom views). **D**) Schematic representation showing the topology of each ArnC protomer, consisting of two transmembrane helices, linked to two juxtamembrane helices (JM1 and JM2), and a large soluble domain consisting of six α-helices and eight β-sheets in a classic GT-A like fold. The C-terminal β8 strand packs with β-strands of the adjacent protomer. **E**) Ribbon representation of one ArnC protomer with the GT-A fold shown in green, and the two juxtamembrane helices in blue. β8’ strand from the adjacent protomer is shown. The numbering of helices and sheets is the same as in (D).

In *E. coli* and *Salmonella enterica*, the most effective modification for reduction of negative membrane charge and development of polymyxin resistance is the attachment of an **aminoarabinose** sugar moiety (4-amino-4-deoxy-L-arabinose or L-Ara4N) to the 1 and 4’ phosphate groups of Lipid A^22,23^. This modification is also critical for development of colistin resistance in *P. aeruginosa*^24^, and *K. pneumoniae*^25^, both included in the ESKAPE pathogen list^26^. L-Ara4N is synthesized by the aminoarabinose biosynthetic pathway, a relay system of eight proteins that act in sequence to synthesize the sugar, associate it with the membrane and transport it to the periplasmic side of bacterial inner membranes^18^. The integral membrane glycosyltransferase ArnC enables the association of the L-Ara4N sugar with bacterial membranes by attaching it to undecaprenyl phosphate (UndP)^27^. ArnC (originally called PmrF) was first identified in *Salmonella* as the product of a gene that disrupted polymyxin resistance when randomly targeted via transposon mutagenesis^28^. It was later demonstrated that ArnC is the glycosyltransferase responsible for attaching the aminoarabinose sugar to the UndP lipid^27^. Only the formylated substrate UDP-L-Ara4FN, produced by the N-terminal domain of ArnA^29,30^, can be converted into the glycolipid product (UndP-L-Ara4FN), whereas the unmodified substrate (UDP-L-Ara4N) cannot be processed^27^ (**Fig. 1A**). ArnC was proposed to resemble dolichol-phosphate mannosyltransferases based on sequence analysis^28,31^. The product of ArnC is deformylated by ArnD^32,33^, flipped to the outer leaflet of inner bacterial membranes by the heteromeric flippase ArnE/F^34^, and finally, processed by the glycosyltransferase ArnT, which transfers the L-Ara4N sugar to Lipid A, enabling resistance to polymyxins and AMPs^22,23,35^.

Glycosyltransferases are classified into families based on amino acid sequence similarity, with 90 distinct families identified to date^36^. Polyprenyl phosphate glycosyltransferases (Pren-P GTs) are a subclass of membrane-bound glycosyltransferases that catalyze the transfer of glycosyl groups from activated sugar donors to a polyprenyl lipid acceptor. This acceptor is then transported across the membrane for use in glycosylation reactions^37,38^. The polyprenyl lipid carrier used by Pren-P GTs varies between organisms, with dolichol phosphate (DolP) used in eukaryotes and archaea, and undecaprenyl phosphate (UndP) used in Gram-negative bacteria^39^. The structure of Pren-P GTs consists of two domains: a cytosolic GT-A-like catalytic domain and a transmembrane domain that anchors the enzyme to the lipid bilayer^38^. The GT-A fold consists of an open twisted β-sheet surrounded by α-helices on both sides, and is reminiscent of two adjoined Rossmann-like folds, typical of nucleotide-binding proteins^36,40^. GT-A enzymes possess a DXD signature in which carboxylates coordinate a divalent cation and/or a ribose^36,40^. Thus far, only two Pren-P GTs have been structurally characterized using X-ray crystallography. GtrB from *Synechocystis sp.* was solved in the presence of nucleotide donor substrate^41^ (PDB code 5EKP), and the dolichol phosphate mannose synthase (DPMS) from *Pyrococcus furiosus* was crystallized in complex with nucleotide, complete donor substrate, and glycolipid product^42^ (PDB codes 5MLZ, 5MM0, 5MM1). Despite the availability of several crystal structures of Pren-P GTs, the catalytic mechanism employed by Pren-P GTs remains poorly understood^38^.

Here we report structures of the Pren-P GT ArnC from the Gram-negative bacterium *Salmonella enterica serovar Typhimurium LT2*, determined by single-particle cryo-electron microscopy (cryo-EM), in its *apo* and partial donor substrate-bound states. By combining structural information with molecular dynamics (MD) simulations, we provide a rationale for substrate binding and propose a hypothesis for the reaction mechanism. Our findings illuminate the structural basis for the catalytic activity of Pren-P GTs and pave the way for developing novel therapeutics targeting polymyxin resistance.

## Results

### Structure determination of ArnC from *S. enterica*

To identify a suitable candidate for structure determination of ArnC, we cloned *arnC* genes from the clinically-relevant species *S. enterica*, *K. pneumoniae*, and *E. coli* with either an N-terminal FLAG-10xHis-TEV cassette or a C-terminal TEV-10xHis cassette using pNYCOMPS vectors^43^. Cloned ArnC orthologs were expressed in an *E. coli* BL21 derivative strain (T7 express lysY/NEB) and extracted with n-Dodecyl-β-D-Maltoside (DDM). Expression was first evaluated in small scale after affinity purification with nickel-affinity resin (**Fig. S1A**). ArnC from *Salmonella enterica* (ArnC*_Se_*) was identified as the most promising candidate, due to the high yield of purified ArnC*_Se_* protein and good stability in detergent. After solubilization with DDM and initial affinity purification, ArnC*_Se_* was reconstituted into lipid-filled nanodiscs for structure determination by cryo-EM. Nanodiscs have been shown to enhance the stability of membrane proteins and provide a more native-like environment for structure determination^44,45^. Based on the size-exclusion chromatography (SEC) profile and SDS page analysis we chose MSP1E3D1 nanodiscs filled with POPG lipid as the optimal condition for ArnC*_Se_* (**Fig. S1B,C**).

For structure determination of the substrate-free (*apo*) state, ArnC*_Se_* embedded in nanodiscs was plunge-frozen in liquid ethane using UltrAuFoil gold grids as support foil to reduce specimen motion^46^. After initial screening, a dataset of 5,552 micrographs was collected using a Talos Arctica cryo-electron microscope equipped with a Gatan K2 direct detector. Processing of this dataset using the workflow described in the methods section resulted in a 2.79 Å reconstruction from ∼184 thousand particles after imposing C4 symmetry (**Fig. S2, Fig. S10,** and **Table 1**). In addition, a second *apo* ArnC*_Se_* dataset of 23,259 micrographs was collected at the National Center for Cryo-EM Access and Training (NCCAT) in New York using a Titan Krios microscope equipped with a Gatan K3 direct detector using a duplicate grid from the same sample preparation. An initial set of over 4 million particles was cleaned up to a final set of ∼490 thousand particles, leading to a 2.74Å reconstruction after imposing C4 symmetry (**Fig. S3, Fig. S10,** and **Table 1**). A direct comparison of the two datasets is discussed in a later section. To generate the initial model, we built into the 2.79 Å substrate-free map reconstructed from the Arctica dataset starting from an AlphaFold generated model of the *E. coli* (K12 strain) ArnC (Uniprot: P77757) from the AlphaFold database^47^. The final model after refinement showed an RMSD of 9.53 Å across 312 Cα atom pairs with the current AlphaFold model of *E. coli* (K12 strain) ArnC in the AlphaFold database (AF-P77757-F1-model_v4) (**Fig. S4A**). The atomic model of ArnC*_Se_* was also refined with the Krios microscope map leading to second model that has 0.67Å RMSD across 1248 Cα atom pairs with the Arctica model. A superposition of the two apo ArnC*_Se_* models is shown in **Fig. S4B**.

**Table 1.**
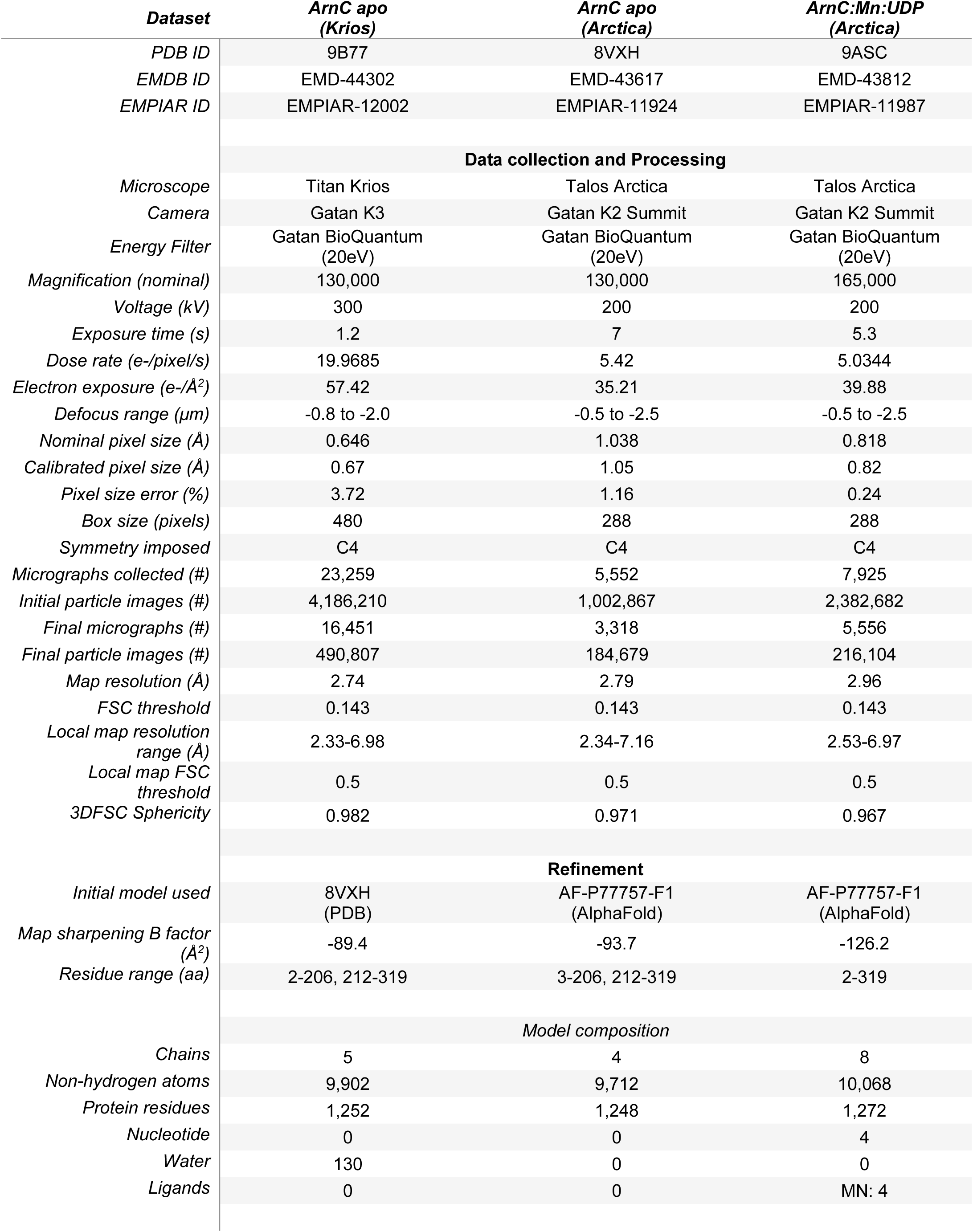
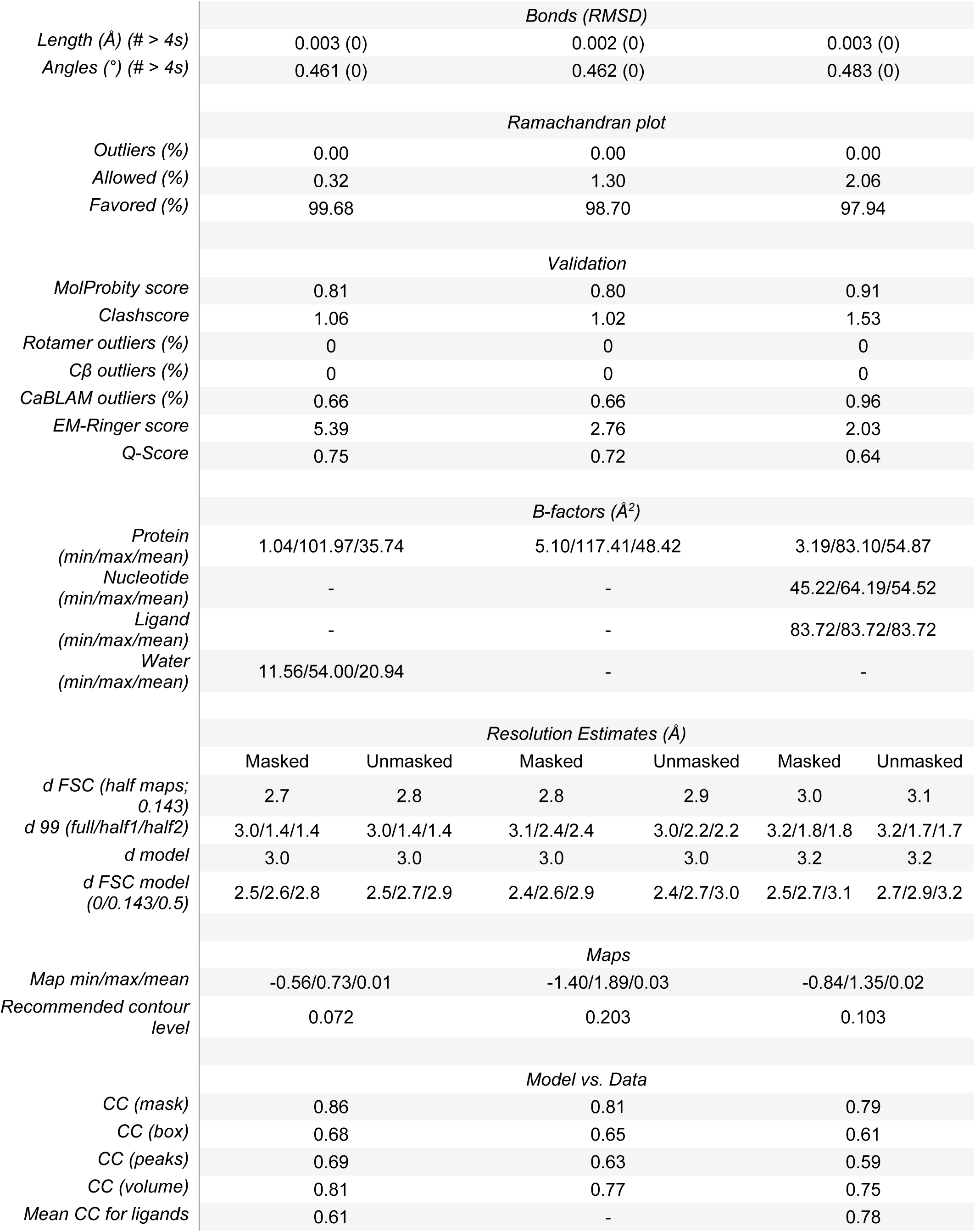
Cryo-EM data collection, refinement, and validation statistics for *apo* ArnC (Krios), *apo* ArnC (Arctica) and ArnC:Mn:UDP complex structures.

| **Dataset** | **ArnC apo**<br>**(Krios)** | **ArnC apo**<br>**(Arctica)** | **ArnC:Mn:UDP**<br>**(Arctica)** |
| --- | --- | --- | --- |
| *PDB ID* | 9B77 | 8VXH | 9ASC |
| *EMDB ID* | EMD-44302 | EMD-43617 | EMD-43812 |
| *EMPIAR ID* | EMPIAR-12002 | EMPIAR-11924 | EMPIAR-11987 |
| **Data collection and Processing** |  |  |  |
| *Microscope* | Titan Krios | Talos Arctica | Talos Arctica |
| *Camera* | Gatan K3 | Gatan K2 Summit | Gatan K2 Summit |
| *Energy Filter* | Gatan BioQuantum<br>(20eV) | Gatan BioQuantum<br>(20eV) | Gatan BioQuantum<br>(20eV) |
| *Magnification (nominal)* | 130,000 | 130,000 | 165,000 |
| *Voltage (kV)* | 300 | 200 | 200 |
| *Exposure time (s)* | 1.2 | 7 | 5.3 |
| *Dose rate (e-/pixel/s)* | 19.9685 | 5.42 | 5.0344 |
| *Electron exposure (e-/Å<sup>2</sup>)* | 57.42 | 35.21 | 39.88 |
| *Defocus range (μm)* | -0.8 to -2.0 | -0.5 to -2.5 | -0.5 to -2.5 |
| *Nominal pixel size (Å)* | 0.646 | 1.038 | 0.818 |
| *Calibrated pixel size (Å)* | 0.67 | 1.05 | 0.82 |
| *Pixel size error (%)* | 3.72 | 1.16 | 0.24 |
| *Box size (pixels)* | 480 | 288 | 288 |
| *Symmetry imposed* | C4 | C4 | C4 |
| *Micrographs collected (#)* | 23,259 | 5,552 | 7,925 |
| *Initial particle images (#)* | 4,186,210 | 1,002,867 | 2,382,682 |
| *Final micrographs (#)* | 16,451 | 3,318 | 5,556 |
| *Final particle images (#)* | 490,807 | 184,679 | 216,104 |
| *Map resolution (Å)* | 2.74 | 2.79 | 2.96 |
| *FSC threshold* | 0.143 | 0.143 | 0.143 |
| *Local map resolution*<br>*range (Å)* | 2.33-6.98 | 2.34-7.16 | 2.53-6.97 |
| *Local map FSC*<br>*threshold* | 0.5 | 0.5 | 0.5 |
| *3DFSC Sphericity* | 0.982 | 0.971 | 0.967 |
| **Refinement** |  |  |  |
| *Initial model used* | 8VXH<br>(PDB) | AF-P77757-F1<br>(AlphaFold) | AF-P77757-F1<br>(AlphaFold) |
| *Map sharpening B factor*<br>*(Å<sup>2</sup>)* | -89.4 | -93.7 | -126.2 |
| *Residue range (aa)* | 2-206, 212-319 | 3-206, 212-319 | 2-319 |
| **Model composition** |  |  |  |
| *Chains* | 5 | 4 | 8 |
| *Non-hydrogen atoms* | 9,902 | 9,712 | 10,068 |
| *Protein residues* | 1,252 | 1,248 | 1,272 |
| *Nucleotide* | 0 | 0 | 4 |
| *Water* | 130 | 0 | 0 |
| *Ligands* | 0 | 0 | MN: 4 |

|  |  |  |  |  |  |  |
| --- | --- | --- | --- | --- | --- | --- |
|  | Bonds (RMSD) |  |  |  |  |  |
| Length (Å) (# > 4s) | 0.003 (0) |  | 0.002 (0) |  | 0.003 (0) |  |
| Angles (°) (# > 4s) | 0.461 (0) |  | 0.462 (0) |  | 0.483 (0) |  |
|  | Ramachandran plot |  |  |  |  |  |
| Outliers (%) | 0.00 |  | 0.00 |  | 0.00 |  |
| Allowed (%) | 0.32 |  | 1.30 |  | 2.06 |  |
| Favored (%) | 99.68 |  | 98.70 |  | 97.94 |  |
|  | Validation |  |  |  |  |  |
| MolProbity score | 0.81 |  | 0.80 |  | 0.91 |  |
| Clashscore | 1.06 |  | 1.02 |  | 1.53 |  |
| Rotamer outliers (%) | 0 |  | 0 |  | 0 |  |
| Cβ outliers (%) | 0 |  | 0 |  | 0 |  |
| CaBLAM outliers (%) | 0.66 |  | 0.66 |  | 0.96 |  |
| EM-Ringer score | 5.39 |  | 2.76 |  | 2.03 |  |
| Q-Score | 0.75 |  | 0.72 |  | 0.64 |  |
|  | B-factors (Å²) |  |  |  |  |  |
| Protein<br>(min/max/mean) | 1.04/101.97/35.74 |  | 5.10/117.41/48.42 |  | 3.19/83.10/54.87 |  |
| Nucleotide<br>(min/max/mean) | - |  | - |  | 45.22/64.19/54.52 |  |
| Ligand<br>(min/max/mean) | - |  | - |  | 83.72/83.72/83.72 |  |
| Water<br>(min/max/mean) | 11.56/54.00/20.94 |  | - |  | - |  |
|  | Resolution Estimates (Å) |  |  |  |  |  |
|  | Masked | Unmasked | Masked | Unmasked | Masked | Unmasked |
| d FSC (half maps;<br>0.143) | 2.7 | 2.8 | 2.8 | 2.9 | 3.0 | 3.1 |
| d 99 (full/half1/half2) | 3.0/1.4/1.4 | 3.0/1.4/1.4 | 3.1/2.4/2.4 | 3.0/2.2/2.2 | 3.2/1.8/1.8 | 3.2/1.7/1.7 |
| d model | 3.0 | 3.0 | 3.0 | 3.0 | 3.2 | 3.2 |
| d FSC model<br>(0/0.143/0.5) | 2.5/2.6/2.8 | 2.5/2.7/2.9 | 2.4/2.6/2.9 | 2.4/2.7/3.0 | 2.5/2.7/3.1 | 2.7/2.9/3.2 |
|  | Maps |  |  |  |  |  |
| Map min/max/mean | -0.56/0.73/0.01 |  | -1.40/1.89/0.03 |  | -0.84/1.35/0.02 |  |
| Recommended contour<br>level | 0.072 |  | 0.203 |  | 0.103 |  |
|  | Model vs. Data |  |  |  |  |  |
| CC (mask) | 0.86 |  | 0.81 |  | 0.79 |  |
| CC (box) | 0.68 |  | 0.65 |  | 0.61 |  |
| CC (peaks) | 0.69 |  | 0.63 |  | 0.59 |  |
| CC (volume) | 0.81 |  | 0.77 |  | 0.75 |  |
| Mean CC for ligands | 0.61 |  | - |  | 0.78 |  |

### The structure of *apo* ArnC

The structure shows that ArnC is a tetramer (**Fig. 1B,C**), with each protomer consisting of an N-terminal cytosolic glycosyltransferase (GT) domain; two juxtamembrane (JM) helices (JM1 and JM2) located adjacent to the transmembrane domain (TMD); two transmembrane helices (TM1 and TM2), which are fully embedded within the nanodisc and anchor the intracellular domains to the membrane; and a short C-terminal β-hairpin (**Fig. 1D,E**). The two transmembrane helices create an eight-helix bundle within the TMD of the tetramer. The short β-hairpin at the C-terminus interacts with the adjacent protomer thus bridging two adjacent subunits within the tetramer. The two amphipathic JM helices are parallel to each other and orientated to expose their hydrophobic faces towards the membrane (**Fig. 1E, Fig. S5C**).

The N-terminal glycosyltransferase GT domain features a classic GT-A fold composed of a seven-stranded β-sheet core flanked by α-helices on both sides. An eighth strand (β8) is contributed by the adjacent protomer and packs against the β3 strand. All eight strands are connected via hydrogen bonds, and two of them (β6 and β8) are antiparallel to the rest. Thus, the ArnC GT-A fold can be described as a Rossmann-like fold consisted of a mixed β-sheet of eight strands ordered in the sequence 83214657 (**Fig. 1D,E**). The conserved signature DXD motif, which is located on the loop between β4 and α4 in ArnC, is known to coordinate the catalytic metal ion and donor substrate in glycosyltransferases that carry a GT-A fold^36^. The ArnC GT-A fold can be further subdivided into a canonical four-stranded Rossmann fold, with four parallel β-strands (β1–β4), two α-helices (α1 and α2) on one side of the sheet, and one α-helix (α3) on the other side, within the N-terminal part of the GT-A fold; and a Rossman-like fold with a three-stranded mixed β-sheet (strands β5–β7) and one α-helix on either side (α4, α5/α6) within the C-terminal part of the GT-A fold (**Fig. 1D,E**).

The catalytic core of the protein between the JM helices and the GT-A domain forms a central cavity (cavity 1) that is lined with conserved charged and polar residues at the periplasmic surface, and hydrophobic residues within the core (**Fig. 2, Fig. S5**). A cluster of highly conserved residues, which includes a number of conserved arginines and the GT-A DXD motif provide a net positive electrostatic charge within this region (**Fig. 2C, Fig. S5**). No visible density was observed within this pocket, verifying that this structure is in its apo state; however amorphous lipid densities could be observed in the inter-protomer grooves of the TM domain, and in particular within a small cavity (cavity 2) formed at the top of the TM domain (**Fig. 2A,B**).

**Figure 2.**
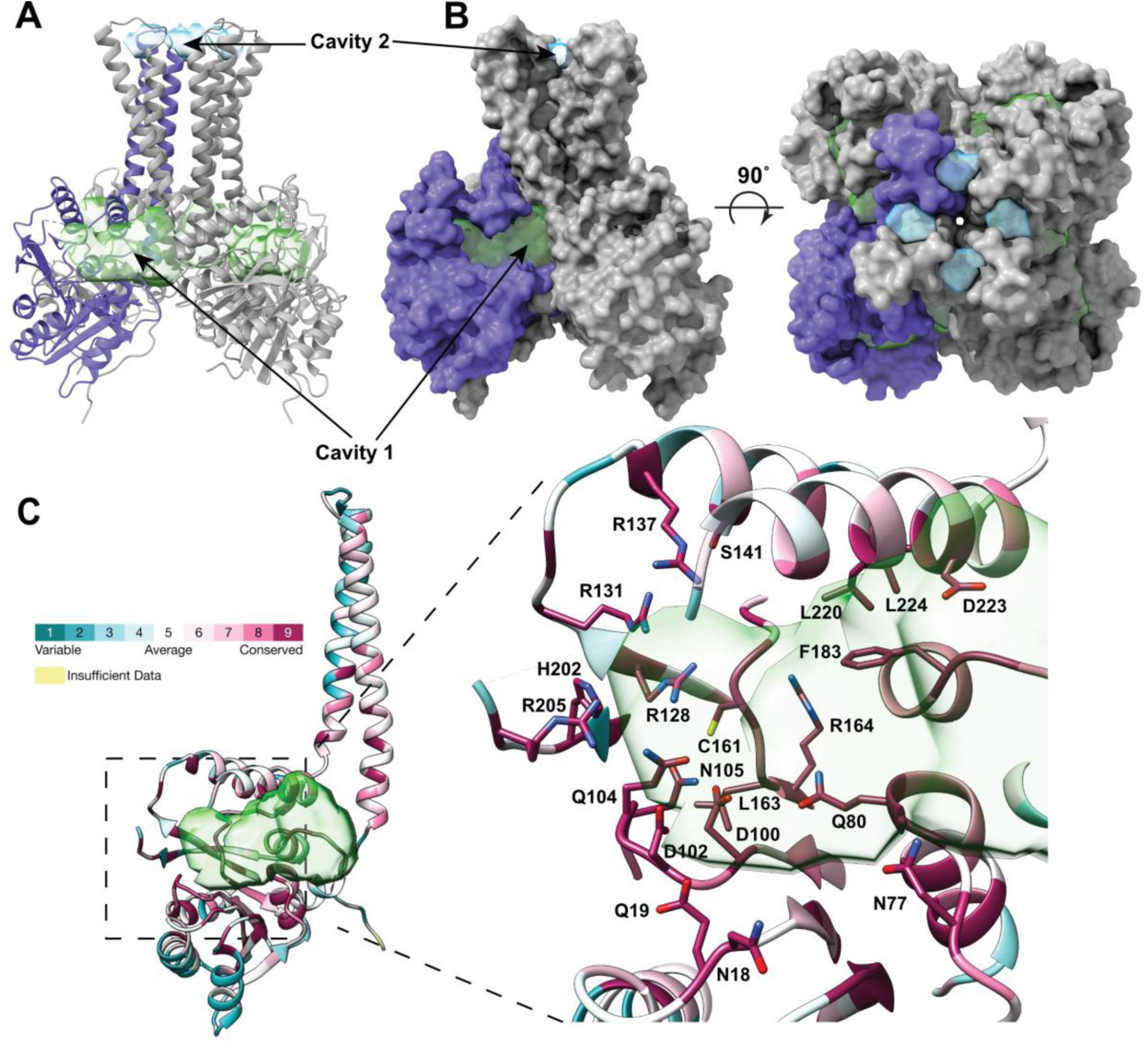
Structural features of ArnC. **A**) Ribbon representation of ArnC showing the volumes of notable cavities. ArnC is colored gray with one protomer colored purple. The volumes are shown as semi-transparent surfaces, with cavity 1 colored green and cavity 2 blue. Volumes were calculated using the Voss Volume Voxelator (3V) server, using probes with 10Å and 2Å radii, corresponding to the outer and the inner probe, respectively. **B**) Spacefill representation of ArnC showing the volumes of notable cavities in two orthogonal views. Coloring is the same as in (A). **C**) Ribbon representation of one ArnC protomer colored by residue conservation from blue (low to no conservation) to purple (absolute conservation). The highly conserved residues are shown in sticks within the binding pocket of cavity 1.

### Comparison of 200kV and 300kV *apo* ArnC datasets

The *apo* ArnC datasets, collected at different sites with different electron microscopes, were refined to 2.74Å and 2.79Å, so we further explored resolution-limiting factors in the datasets collected on the Arctica and the Krios microscopes. To understand factors governing resolution from the two *apo* ArnC datasets, we first repeated the final 3D reconstruction with subsets of various sizes, randomly sampling particles present in the full datasets. At 180,000 particles (the largest subset contained in both datasets), ArnC was resolved to 2.81Å at 300 kV and 2.80Å at 200 kV. At all particle counts (from 2,500 to 180,000 particles), the difference in resolution (at FSC=0.143) was small between datasets acquired at the two sites, averaging 0.04Å better at 300kV (**Fig. S6A,C,D**). The most reasonable explanations for this difference include: effect of higher magnification (0.646Å/px vs. 1.038Å/px), effect of detector performance (Gatan K3 vs. Gatan K2 direct electron detector), effect of microscope model and voltage (300kV Titan Krios vs. 200 kV Talos Arctica), effect of the experimenter/site, or other uncontrolled variation between experiments (e.g. intrabatch variation in grid quality, environmental interference, day-to-day variation in etc.). The difference of 0.04Å resolution did not lead to meaningful differences in the biological interpretation of the macromolecular structure (**Fig. S4B**). We conclude that biochemical and/or sample preparation factors are more significant limiting factors for ArnC structure determination than electron microscope voltage.

In theory, the number of particles required to reach certain resolution is proportional to *e^B^*^/2d²^, where *B* is the temperature factor and *d* is the resolution^48^, and *B* will be improved by superior microscope/detector quality. Varying particle number by reconstructing random subsets allows us to assess the conformance of the actual data to this prediction. In both datasets, resolution improves with increasing particle count in line with theory^48^ and the B-factor plot is linear for all particle counts (**Fig. S6B**). Holding particle alignment fixed, the spectral signal-to-noise ratio (SSNR) of the reconstruction is simply the product of the per-particle SSNR with the particle count^49^. Indeed, where the FSC>0.143, this relationship holds at all particle counts tested (up to 490,000 particles, or 1.96×10^6^ asymmetric units, for the Krios dataset); this is demonstrated in the “universal plot” of ln(SSNR/*n*) vs. spatial frequency^49^ by the fact that the lines for each particle count are indistinguishable at resolutions >3Å (**Fig. S6E,F**). The SSNR has features in the same locations as the radially-averaged structure factor of the reconstructed map, but convolved with a dampening factor representing resolution-dependent attenuation of information transfer. Extrapolating from the calculated B-factor, reaching 2.0Å would take one millionfold more particles, whereas such resolution is readily attained under these conditions with 490,000 apoferritin particles. Although ArnC resolution did not reach a resolution cutoff and may continue to improve with particle count as *n* ∝ *e^B^*^/2d²^, the B-factor (which was similar between microscopes/cameras/voltages) implies that resolution would be practically limited to ∼2.5Å unless biochemical or grid conditions were substantively different. This differs from the case of an 85kDa CDK-activating kinase, where voltage effects were the most likely driver of a 0.3Å resolution difference^50^, and suggests that resolution gains with use of higher microscope voltage will be specimen-dependent.

### Comparison with existing Pren-P GT structures

In addition to the structures reported here, the structures of two other Pren-P GTs have been solved previously by X-ray crystallography. In this section we provide a comparison between ArnC and the two existing Pren-P GT structures. GtrB is a glycosyltransferase that catalyzes the transfer of glucose, from UDP-Glucose to the undecaprenyl phosphate (UndP) lipid carrier^41^. The structure of GtrB from Synechocystis species^41^ (PDB code 5EKP) has been determined in a nucleotide bound state at 3.2Å resolution. Similar to ArnC, GtrB is a tetramer with two TM helices and a classic GT-A domain at the N-terminus (**Fig. S7A,B**). ArnC and GtrB have overall similar topology. Yet, the two TM helices in GtrB assume different positions relative to the catalytic domain compared to ArnC (**Fig. S7C**). The crystal structure shows that GtrB coordinates a UDP nucleotide within a GT-A domain pocket utilizing a magnesium metal to coordinate the phosphate groups of the donor. This pocket corresponds to cavity 2 in ArnC (**Fig. 2**). GtrB utilizes a DXD motif (_94_DADLQD_99_) to coordinate the magnesium metal using D96 and Q98, which is similar to the DXD motif of ArnC (_100_DADLQN_105_). In addition, the loop containing the DXD motif appears to be in a similar conformation in the two enzymes (**Fig. S7D,E**).

Dolichyl phosphate mannose synthase (DPMS) catalyzes the transfer of mannose from the donor GDP-mannose to dolichol phosphate (DolP) to produce DolP-mannose (DolP-Man)^42^. The structure of DPMS from *Pyrococcus furiosus* (*Pf*DPMS) has been determined by X-ray crystallography in complex with nucleotide GDP, full donor substrate (GDP-Man), and glycolipid product (PDB code 5MLZ, 5MM0 and 5MM1)^42^. Unlike ArnC, *Pf*DPMS is a monomer in the crystal state (or perhaps a dimer in its physiological state), and has four TM helices, arranged as two α-helical hairpins (**Fig. S8A,B**)^42^. Due to the differences of the TM helices, a structural alignment of *Pf*DPMS with an ArnC protomer is only meaningful for the GT-A domain (**Fig. S8C**). When only the GT-A domain of ArnC and *Pf*DPMS are aligned, they superimpose well (14.7Å RMSD across 388 Cα atom pairs). Based on the *Pf*DPMS structure in the product-bound state (PDB 5MM1), the JM helices enable the threading of the acceptor lipid DolP within the active site of the GT-A domain. A similar function for the JM helices of GtrB was hypothesized based on the GtrB crystal structure^41^. ArnC exhibits similar JM helices that may subserve the same role. This will be examined in a later section. Finally, like GtrB, PfDPMS utilizes a DXD motif (_89_DADLQH_94_) to coordinate a Mg^2+^ or Mn^2+^ metal and the donor nucleotide (**Fig. S8D,E**).

### The structure of UDP-bound ArnC

Given the similarity of ArnC to GtrB and *Pf*DPMS, and the presence of a DXD motif, ArnC likely coordinates its nucleotide-sugar donor substrate via a metal ion. Our first goal was to confirm metal binding and identify the optimal metal ion for structural studies of ArnC in a nucleotide-bound state. Thus, we used Microscale Thermophoresis (MST) to characterize the interaction between ArnC and the nucleotide UDP either in the presence of Mg^2+^ or Mn^2+^ metal ions, or in the absence of metal ions (**Fig. 3A,B**). MST is a biophysical technique that enables quantification of subtle changes in fluorescence in response to ligand binding, while the target protein is subjected to a thermal gradient^51^. UDP was titrated into ArnC in the presence of either 1mM MnCl_2_, 1mM MgCl_2_, 1mM EDTA to chelate any endogenously-bound metals, or without any exogenous metal added (**Fig. 3B**). A clear difference was observed when titrating UDP in the presence of MnCl_2_ compared to the other three conditions. Calculated Kd values based on the fitted curves were 8.5 µM in the presence of MnCl_2_, 320 µM in the presence of MgCl_2_, 8.5 mM in the presence of EDTA, and 591 µM without addition of exogenous metals. Only the curve fitted to the titration in the presence of MnCl_2_ is a high confidence fit, as judged by the reduced Chi square value. An additional caveat for this experiment is that we were unable to strip the protein of endogenous metals before the MST measurements because extensive treatment with EDTA caused the protein to crash out of solution. Thus, Kd values presented here should only be considered in relative terms rather than absolute, since it is very likely that only a portion of the metal binding sites of ArnC were available to bind the exogenous metals added. Yet, this experiment shows clearly that Mn^2+^ enables higher affinity of ArnC towards the UDP nucleotide, suggesting that it maybe the preferred metal for the ArnC metal-binding site(s).

**Figure 3.**
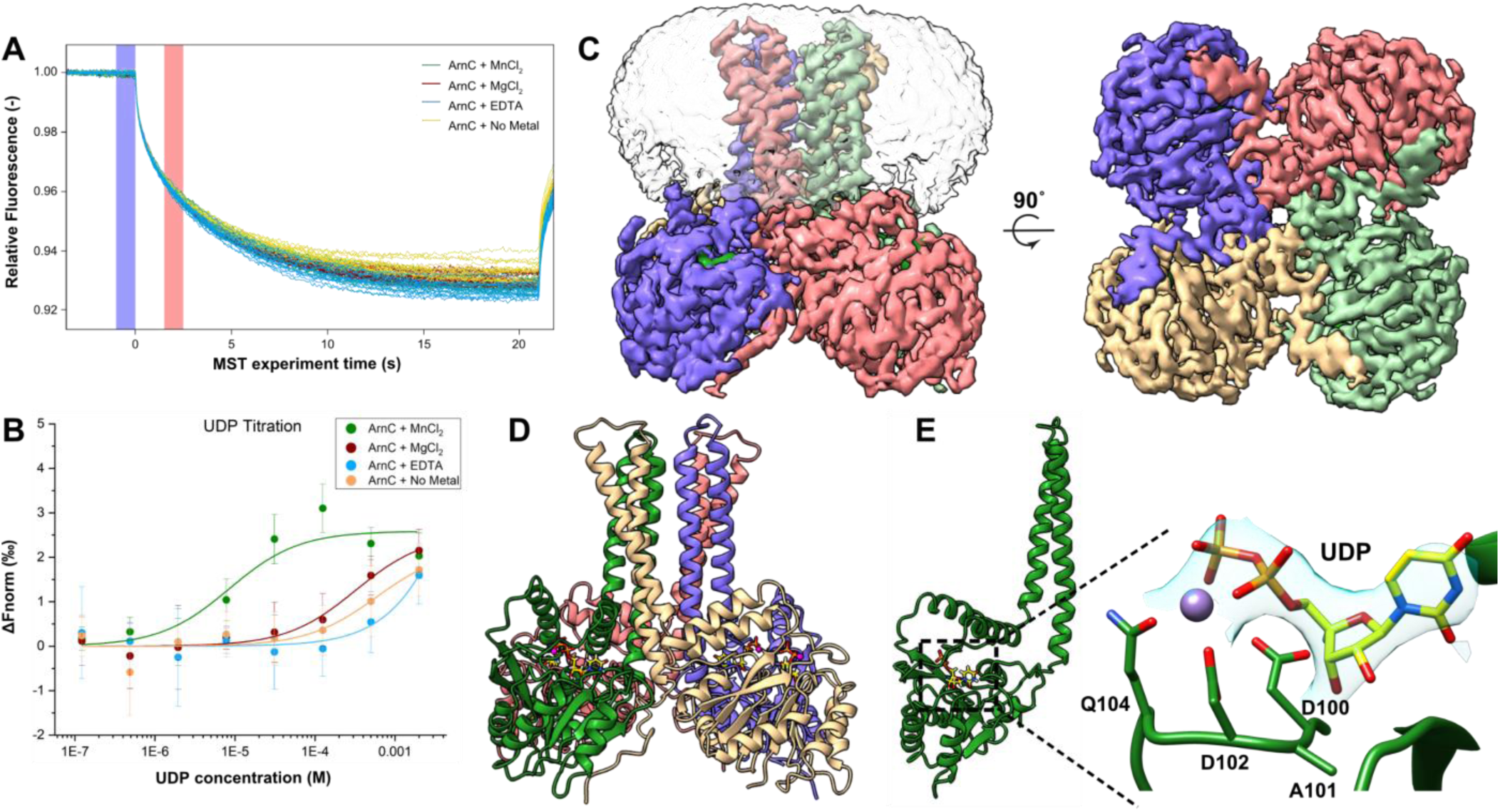
The cryo-EM structure of the UDP-bound state of ArnC. **A**) Overlay of representative MST trace for all four experimental groups, showing the cold region (−1 to 0s) and hot region (1.5 to 2.5s) used for analysis of the thermophoresis traces. Traces are colored according to the inset; MnCl_2_ (Green), MgCl_2_ (Red), No added metal (Yellow) and EDTA (Blue). **B**) Scatter plot showing the composite of three independent experiments for the titration of UDP under the four different experimental conditions. The Kd values calculated based on the fitted curves are: 8.48±6.83μM with 1mM MnCl_2_, 323.44±106.12μM with 1mM MgCL_2_, 8.55±32.65mM with 1mM EDTA, and 0.59±0.57mM with no exogenous metal added. The presence of Mn^2+^ leads to a pronounced increase in the binding affinity of ArnC for the UDP ligand. Error bars represent standard deviation (SD). **C**) Cryo-EM density map of the ArnC tetramer in the presence of Mn^2+^ and UDP in two orthogonal views. Coloring is per protomer, and the semi-transparent volume represents the nanodisc density observed in low contour levels of the reconstruction. **D**) The cryo-EM structure of ArnC bound to Mn^2+^ and UDP. The coloring is per protomer, with UDP in yellow, the phosphates of UDP in orange, and the Mn^2+^ ion represented as a purple sphere. **E**) A single protomer of ArnC bound to Mn^2+^ and UDP is shown. The magnified detail shows the characteristic _100_DXD_102_ motif found within the GT-A domain that facilitates coordination of the metal and UDP nucleotide. The cryo-EM density around UDP is shown as a semi-transparent surface. Coloring is the same as in (D).

To prepare UDP-bound ArnC for cryo-EM analysis, 2mM UDP and 1mM MnCl_2_ were added to the purified protein prior to vitrification. 7,925 movies of UDP-bound ArnC were collected using a 200 kV Talos Arctica microscope equipped with a Gatan K2 direct detector. Processing of this dataset using the workflow described in the methods section resulted in a 2.96 Å reconstruction from 216 thousand particles after imposing C4 symmetry (**Fig. S9, Fig. S10,** and **Table 1**). Analysis of the UDP-bound ArnC map shows an overall fold similar to the *apo* ArnC structure (**Fig. 3C,D**). Within the binding cavity of the GT-A domain we observed clear non-protein density, which we modeled as UDP coordinated by the Mn^2+^ metal ion (**Fig. 3E** and **Fig. S10**). The UDP nucleotide binds in a shallow cleft on the surface of the GT-A domain, adjacent to the DXD motif, with D102 and Q104 clearly coordinating the Mn^2+^ metal ion, which in turn coordinates the diphosphate of the UDP (**Fig. 3E**). The UDP molecule is also coordinated via hydrogen bonds with the sidechains of S210, R205, N77, D48 (**Fig. S11A**), and backbone atoms of V16, A101 and K211 (not shown). A comparison of this binding mode with nucleotides bound to the other Pren-P GT structures shows that coordination and localization of the nucleotide within the GT-A domain is similar between ArnC, GtrB and *Pf*DPMS (**Fig. S11A**).

All structures of Pren-P GTs to date have been determined in a nucleotide-bound state. Since we have now obtained structures of a true *apo* state of ArnC and a nucleotide-bound state, we further compared the structures of the two states by aligning them (**Fig. 4A**). After aligning the two structures on their transmembrane domain (aa 230-305), we detected a significant conformational transition that occurs when UDP binds to its binding cleft (**Fig. 4A,B**). The overall fold of the GT-A domain remains largely unchanged between the two states, but in the UDP-bound state the GT-A domain comes closer to the JM helices (and the membrane). There is some movement at the JM helices (more in JM1 than JM2), but the major conformational changes occur in the GT-A domain, which moves like a “pendulum” upwards (**Fig. 4B**). The GT-A domain rotates upwards by 13.7° and different parts of the domain undergo a translation ranging from 2.7 to 7.7Å. Moreover, the linker between strand β7 and JM2 becomes more ordered and can be fully modelled in the UDP-bound structure (aa 207-211 are missing from the *apo* structure). The conformational transition observed here is a typical clamshell motion to restrict the nucleotide within the binding cleft, and has been observed in cases of other domains binding to nucleotides (e.g. UDP-glucose 6-Dehydrogenase UGDH, UDP-MurNc-tripeptide ligase MurE, nucleoside triphosphate diphosphorylase NTPDase1)^52–54^. This transition and the concomitant binding of the nucleotide substrate likely represents the faster kinetic component of the enzymatic cycle compared to binding of the lipidic acceptor UndP into the binding site.

**Figure 4.**
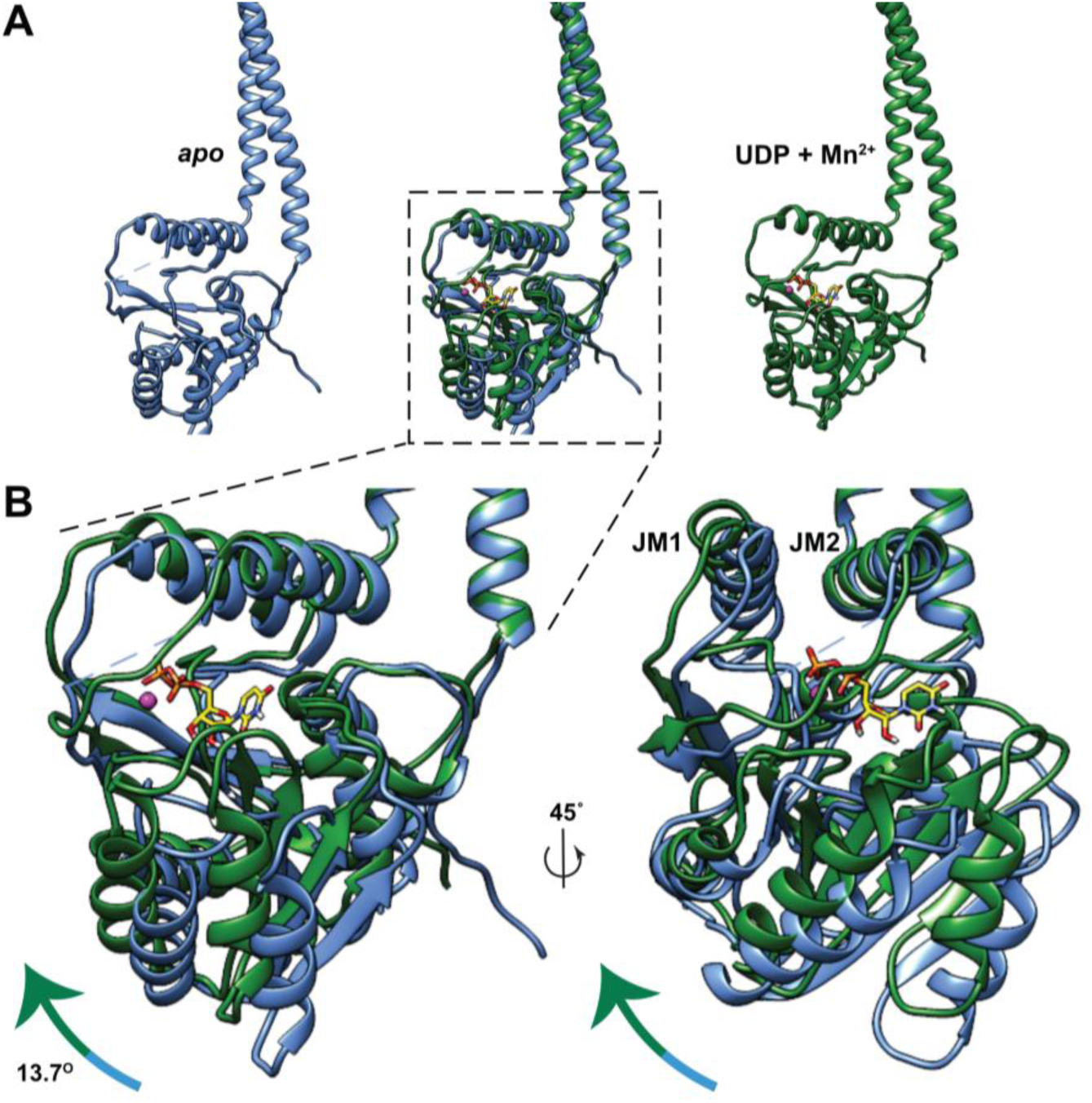
ArnC undergoes a conformational transition upon nucleotide binding. **A**) Superposition of apo ArnC (blue) and UDP-bound ArnC (green) structures, with a single protomer from each structure aligned at the transmembrane helices (aa 230-305). **B**) Detail from (A). Two views of the superposition rotated by 45° are presented. An upward movement is observed in the GT-A domain when the nucleotide is bound, with the domain rotating upward by 13.7^°^, and different parts of the domain showing a translation between 2.7-7.7Å. The Mn^2+^ ion is shown as a pink sphere, and the UDP nucleotide is shown in stick representation and yellow color, with the phosphates shown in orange color.

### Putative UndP-binding motif of ArnC

In the crystal structure of *Pf*DPMS bound to dolichol phosphate mannose (DolP-Man) (PDB 5MM1), the lipid DolP can be seen threading between the juxtamembrane helices to reach the active site of the GT-A domain^42^ (**Fig. S14A**). A similar function for the JM helices of GtrB was hypothesized based on the GtrB crystal structure^41^. We propose that in ArnC the JM helices act in a similar manner to facilitate the entry of the acceptor substrate undecaprenyl phosphate (UndP) in the active site of ArnC. In addition, several non-protein densities likely corresponding to annular lipids were observed adjacent to TMD helices and especially within cavity 2 and at the base of the TMD close to the JM helices (**Fig. 5A**). To investigate these densities and the possible binding interface of UndP, we utilized LipIDens, a pipeline for molecular dynamics (MD) simulation-assisted interpretation of lipid and lipid-like densities in cryo-EM structures^55^. To assign the most likely identity of the additional lipid-like densities in the cryo-EM map we performed coarse grade (CG) simulations of ArnC initiated from the apo conformation. Cardiolipin preferentially bound to a groove on the periplasmic TMD face with a residence time ∼0.4-0.5 μs (**Fig. 5A**). The top half of the transmembrane domain contains only two positively-charged or polar residues, R258 and Q264, both located near cavity 2. This may account for the reduced residence time compared with high affinity cardiolipin sites across other prokaryotic membrane proteins^56^. Surprisingly, we observe spontaneous and stable binding of UndP (∼8 μs) or a single tail of POPE (∼10 μs) within the GT-A domain (**Fig. 5A**) whereby the phosphate group of UndP extends beyond the plane of the bilayer phosphate beads in the top ranked CG binding pose (**Fig. 5B**).

**Figure 5.**
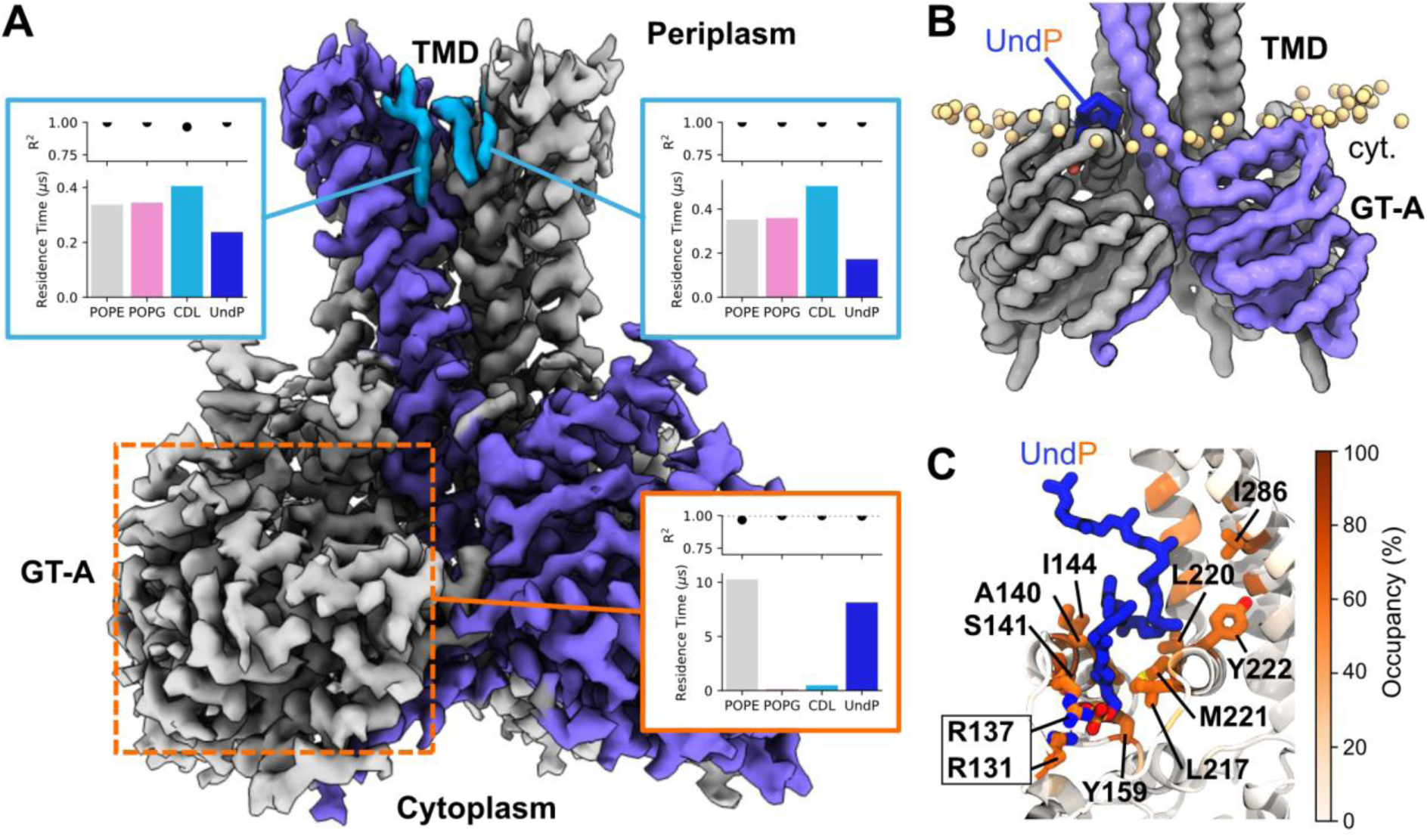
UndP and lipid interactions with ArnC. **A**) Additional densities surrounding the apo ArnC TMD, colored according to preferential lipid interactions in coarse-grained (CG) MD simulations (10 x 15 μs) embedded within a bilayer contained POPE (64%), POPG (18%), cardiolipin (CDL) (9%) and UndP (9%). Plots indicate the relative residence times of lipids binding to the sites, as calculated using PyLipID. We also observe spontaneous binding of UndP within the GT-A domain in CG simulations (boxed in orange) and indicated in (B) and (C). **B**) Top ranked CG binding pose for UndP within the GT-A domain. UndP is colored blue and the UndP phosphate bead is shown in orange with respect to the position of all lipid phosphate beads within the bilayer (yellow). **C**) UndP pose at the end of atomistic MD simulations (3 x 100 ns) initiated by back-mapping from the top ranked CG UndP pose in (B). Residues with interaction occupancies > 85% across all replicates are shown as sticks and labelled.

To further assess the binding of UndP within the GT-A domain, we performed atomistic MD simulations initiated from the top ranked CG pose. UndP was stably coordinated within the GT-A domain. The tail inserts between JM1 and JM2, forming hydrophobic interactions with small aliphatic sidechains (Ala, Leu, Ile). In contrast, the phosphate headgroup of UndP is coordinated by R131 and R137 (**Fig. 5C**). This binding pose is in broad agreement with that predicted for GtrB^41^, and observed within the structure of *Pf*DPMS bound to DolP-Man^42^. Furthermore, we compared the position of UndP in the GT-A domain (as predicted from simulations) with the UDP binding pose in the UDP-bound ArnC cryo-EM structure (**Fig. S11B**). Both UndP and UDP can be coordinated within the GT-A domain without coordination clashes. We also observed a rearrangement of a flexible loop (residues 201-208) that is stabilized by the presence of UndP compared to the *apo* conformation (**Fig. S11C**). This rearrangement leads R128 and R205 to face towards the UDP binding site at the end of the simulations. R205 can be seen in the UDP-bound ArnC cryo-EM structure coordinating the UDP nucleotide (**Fig. S11A**), and the flexible loop that “covers” the active site in each GT-A domain becomes fully ordered in the presence of the UDP nucleotide (aa 207-211 are only visible in the UDP-bound ArnC cryo-EM structure). Thus, the flexible loop appears to be stabilized by binding of either UndP or the UDP nucleotide, based on the simulations and the UDP-bound cryo-EM structure. Residues R131, R137, and perhaps R128 appear to participate in coordination of UndP, while R205 coordinates the incoming nucleotide (together with the metal ion and the additional residues described in the previous section).

### Molecular modelling and atomistic simulations of substrate-bound ArnC

In order to better describe the catalytically-relevant residues of ArnC, we proposed three models for the enzyme in complex with its full substrates, UndP and UDP-L-Ara4FN (UDPA) (**Fig. 6A**). For the first two models (State 1 and State 2), the lipid was positioned based on the binding observed in the coarse-grained simulation, with the phosphate of UndP coordinated by R128 and R137. For State 1, the Ara4FN group of UDP-L-Ara4FN was placed in a groove between the JM2-*β*7 and JM1-*β*5 loops with the C1 atom within a hydrogen-bonding distance of the UndP phosphate group (State1 – **Fig. 6A**). For the second model, the orientation of the Ara4FN group was based on the structure of PfDPMS in complex with GDP-mannose (PDB 5MM0) (State 2 – **Fig. 6A**). To allow the reaction to occur, a third model was constructed in which the lipid was positioned deeper inside the active site, below the sugar, with the phosphate group within a hydrogen-bonding distance of the C1 atom of the Ara4FN group (State 3 – **Fig. 6A**). UDP-L-Ara4N exhibits increased flexibility within the active site when initially placed in state 1 (**Fig. 6B,C**– bottom left panel). For all states the UDP portion of the UDP-L-Ara4N interacts with ArnC in a similar way, through several possible hydrogen bonds involving residues P15, Y17, E19, D48, A101 and R205. The diphosphate-binding site is formed at the canonical DXD motif, with hydrogen bonds between the diphosphate moiety of UDP and R205 also observed (**Fig. 6D** and **Fig. S12**). While the UDP portion of UDP-L-Ara4FN behaves similarly for all models, the Ara4FN group is much more flexible within the active site for the state 1. Interestingly, for this state, the sugar group rotates in the opposite direction of its initial configuration, adopting a binding pose similar to the one seen in the state 2 and 3 at the end of the simulation (**Fig. 6B**). For all states, hydrogen bonds between R137 and Ara4FN group were observed suggesting that R137 may also act in coordination of the sugar and stabilization of the substrate during catalysis. The high stability of UDPA within its binding site observed for states 2 and 3 suggest that the second proposed binding mode is more favorable.

**Figure 6.**
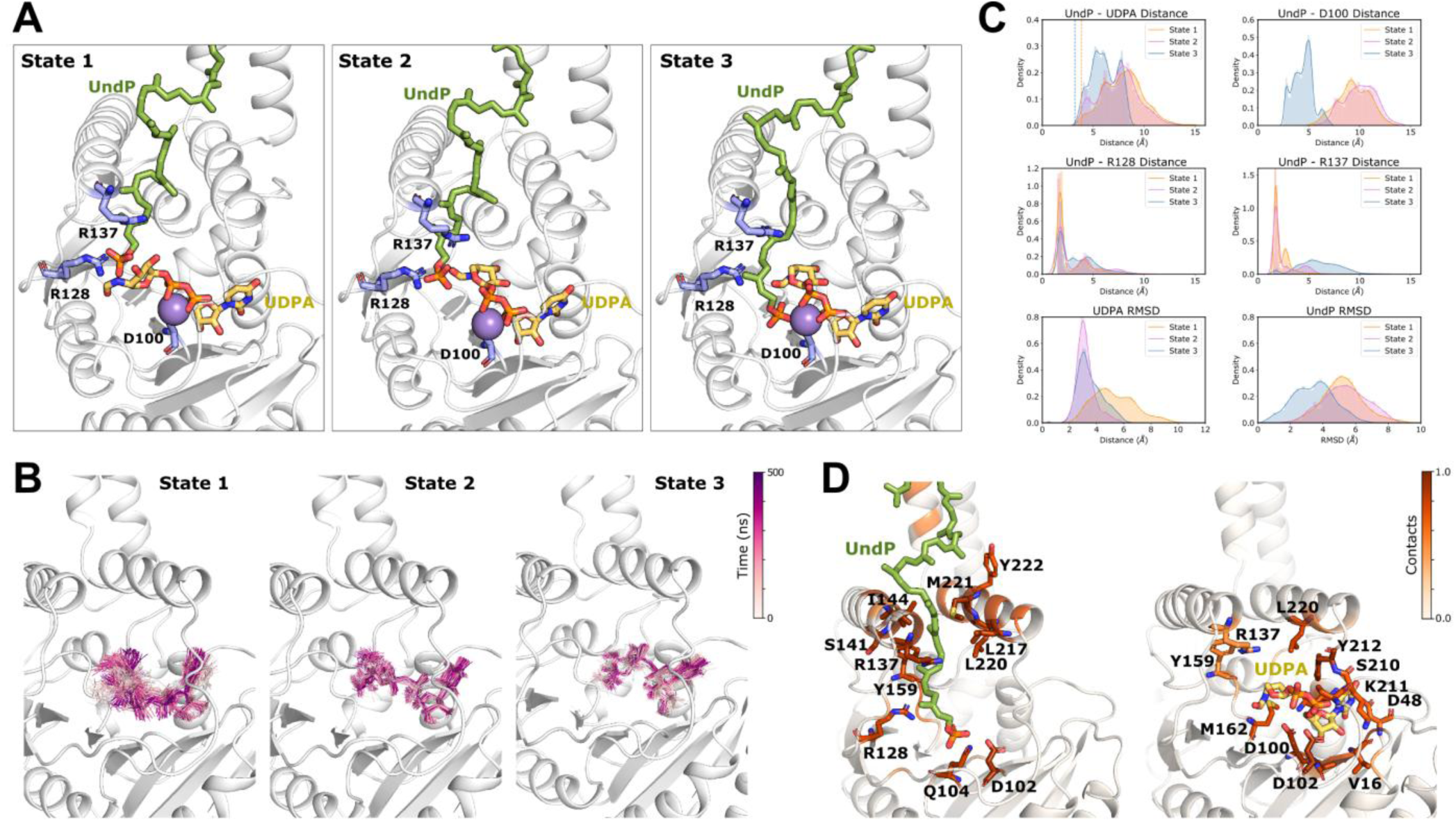
Atomistic simulations of ArnC with both substrates. **A**) Models for the ArnC in complex with its substrates, UndP and UDP-L-Ara4FN: two conformations for UDP-L-Ara4FN and the UndP were modelled. In the first model the lipid was placed in the binding site predicted in the binding coarse grained simulation, with its phosphate group coordinated by the conserved arginines R128 and R137. The Ara4FN group was positioned in an open groove within within hydrogen-bonding distance from the phosphate of UndP (State 1). For the second model, the position of the sugar group was based on the structure of *Pf*DPMS in complex with GDP-mannose (PDB ID: 5MM0) while the coordinates for UndP was kept the same (State 2). For the third model, the lipid was positioned deeper inside the catalytic site, below to the sugar (State 3). **B**) Superposition of multiple frames showing the flexibility of UDP-L-Ara4FN within the ArnC active site for states 1,2 and 3. **C**) UndP-UDPA distance distributions. The distance between UndP and UDPA was defined as the distance between the protonated oxygen of the phosphate group and the arabinose C1 atom (top left panel). UndP-D100 distance distributions (top right panel). UndP-R128 distance distributions (middle left panel). UndP-R137 distance distributions (middle right panel). The distances between UndP and D100, R128 or R137 were defined as the minimum distance between the phosphate group and each of the residues sidechains. UDPA (bottom left panel) and UndP RMSD (bottom right). The RMSDs were calculated using the protein as reference. To calculate the UndP RMSD only the phosphate group atoms and the first carbon were considered, due the high flexibility of the lipid tail. **D**) Contacts between ArnC and UndP or UDPA for state 2. Coloring is the same as in (A).

To evaluate the viability of the proposed reaction models we measured the distance between UndP and UDP-L-Ara4N over the simulation (**Fig. 6B** – top left panel). This distance was defined as the distance between the protonated oxygen of the phosphate group and the arabinose C1 atom. Even though for states 1 and 3 the UndP phosphate group was initially placed within 3.5Å of the C1 atom, the higher distances sampled and the wider distance distribution for state 1 compared to state 3 suggest that the third model is most likely. The increased distance between UndP and UDPA observed in state 1 simulations is mainly due to a rearrangement in the lipid position, which is displaced toward the exit of the active site. In this position, the phosphate group retains its coordination by the R137 and R128 residues (**Fig. 6C**). While the phosphate group is more dynamic in state 1 (**Fig. 6C** – bottom right panel), for state3, its coordination by Mg^2+^ and the R128 residue stabilizes the phosphate in the proximity of the Ara4FN group over the duration of the simulation. Moreover, for this state, the short distance between the UndP phosphate group and D100 (the first D in the DXD motif) suggests that this residue may act in the deprotonation of the phosphate group to allow the reaction to proceed (**Fig. 6C** – top right panel).

Even though the binding coarse-grained simulation and the atomistic simulations of state 2 suggest a somewhat different coordination for the UndP phosphate group, the CG simulations were conducted in the absence of UDPA and the divalent cation. Therefore, the CG and atomistic simulation are likely capturing different aspects of the catalytic cycle of the enzyme. The lipid would initially access the active site via the mechanism proposed by the CG simulation and move deeper inside the catalytic site when in the presence of UDPA and Mg^2+^. Moreover, the similar behavior of UDPA observed for states 2 and 3 regardless of the position of the lipid, indicates that the presence of the lipid is not required for UDPA binding.

### Mechanism of catalysis

The GT-A domain of ArnC is similar to the GT-A domain of Pren-P GTs GtrB and *Pf*DPMS, thus, comparison of the putative catalytic residues that operate in these three enzymes can help us better define the mechanism of catalysis for transfer of a sugar moiety from a nucleotide donor (UDP or GDP) to a lipid phosphate acceptor (UndP or DolP) in Pren-P GT enzymes. Based on the crystal structures of *Pf*DPMS bound to full donor substrate GDP-mannose (PDB 5MM0) and to product DolP-mannose (PDB 5MM1), it was proposed that dolichol phosphate is positioned within the active site by S135, R117 and R131 for catalysis^42^ (**Fig. S13A, S14A**). The acceptor phosphate of DolP, being in the dianionic form, is then suggested to perform a direct nucleophilic attack on the C1 atom of the mannose sugar^42^. All three of these residues are conserved in ArnC as S141, R128, and R137, which also has been seen interacting with the acceptor UndP in our CG and atomistic simulations (**Fig. 5A, 6A, S12**). In addition, R205 in ArnC which is fully conserved in both other enzymes (R202 in *Pf*DPMS) participates in the coordination of the donor phosphate group (**Fig. S13A,B**). The crystal structure of GtrB (PDB 5EKP) is missing densities for both β5-JM1 and β7-JM2 loops, has JM1 missing in some protomers, and missing sidechains for some critical residues, making comparison with ArnC more difficult. Despite that, R122 in GtrB likely corresponds with R117 in *Pf*DPMS and R128 in ArnC (**Fig. S13A**), and K132 is the equivalent residue to R131 in *Pf*DPMS and R137 in ArnC (**Fig. S13B**). R122 in GtrB has been proposed to coordinate the acceptor phosphate^41^, similarly to the equivalent residues in *Pf*DMPS and ArnC. For GtrB, D157 was proposed to act as a catalytic acid that will donate a proton to the distal phosphate of the donor, after the acceptor phosphate of UndP in its dianionic form performs a direct nucleophilic attack on the C1 of the glucose sugar^41^ (**Fig. S13A**). The equivalent residue in *Pf*DPMS is a Gly and in ArnC a Met (**Fig. S13B**). Thus, this mechanism is unlikely to operate in either *Pf*DPMS or ArnC.

In both *Pf*DPMS and GtrB, it has been proposed that the acceptor phosphate in its dianionic form performs a direct nucleophilic attack to the C1 of the sugar bound to the nucleotide donor to create the glycosidic bond of the product. Yet, phosphomonoesters have two pKas, with one expected to be in the physiological pH range^57^. The proportion of polyprenol phosphate lipids that are in their dianionic form at physiological pH has not been determined as far as we know. The secondary pKa (pKa2) of phosphatidic acid (PA) and lysophosphatidic acid (LPA), which also carry a single phosphate connected to a glycerol and two or one acyl chain, respectively, has been measured to be between 6.8-7.5 for LPA and 6.9-7.9 for PA, and vary based on the lipid composition of the bilayer^57^. In monolayers, PA has been determined to have an even higher pKa2 value (10.5-11.5), which is also dependent on salt concentration^58^. Thus, it is very likely that at physiological pH a significant proportion of polyprenol phosphate acceptor lipids are in their monoanionic form, which would imply that simple coordination of UndP (or DolP) within the active site would not be sufficient for catalysis. We have observed that in all three cases of Pren-P GT enzymes, the first aspartate of the **D**XD motif (D89 in *Pf*DPMS, D94 in GtrB and D100 in ArnC) does not participate in the coordination of the catalytic metal ion and is available to act as a catalytic base. Our atomistic simulations have shown that the acceptor phosphate of UndP is coordinated in the proximity of D100 (**Fig. 6C, Fig. S14C**). In addition, in *Pf*DPMS it has been shown that a D89A mutation leads to reduction of catalytic activity similar to the S135A mutation, while the second aspartate of the DX**D** motif when mutated (D91A), does not lead to as severe reduction of catalytic activity^42^. This argues that D89 may also operate as a catalytic base in *Pf*DPMS. In **Fig. 7B** we propose a catalytic mechanism for ArnC, which shows that a monoanionic form of UndP is coordinated in the active site via interactions with R128 and R137. D100 abstracts a proton from the acceptor phosphate and activates it. Subsequently, the acceptor phosphate performs an S_N_2-like nucleophilic attack to the C1 atom of the L-Ara4FN sugar to create UndP-L-Ara4FN with inversion of the anomeric carbon. This reaction mechanism likely operates on all Pre-P GT enzymes, as the DXD motif is conserved throughout the family.

**Figure 7.**
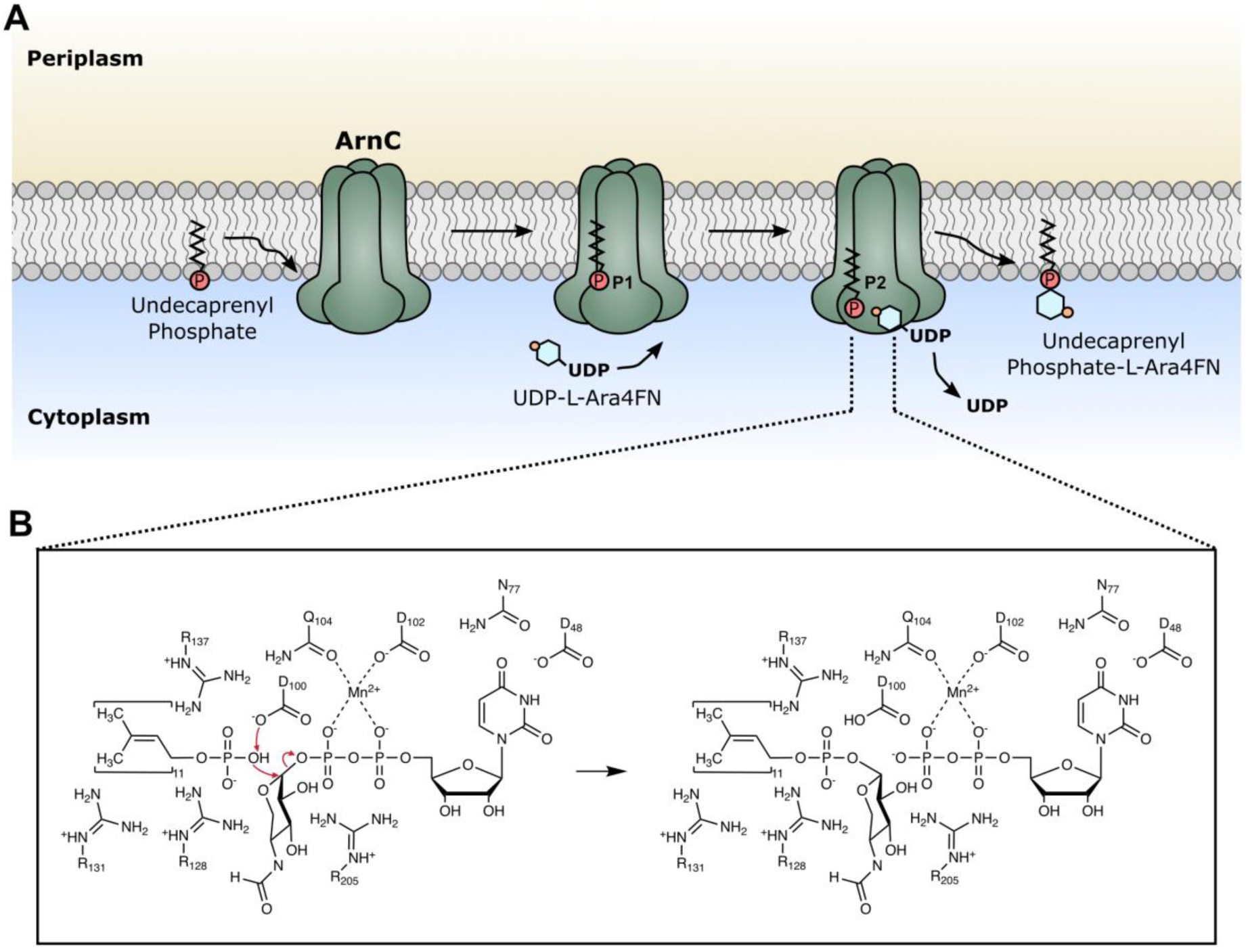
The catalytic mechanism of ArnC. **A**) Schematic representation of the catalytic cycle of ArnC. The acceptor lipid undecaprenyl phosphate (UndP) enters the enzyme (by threading via the JM helices into the GT-A domain), and is coordinated in position 1 (P1), the “standby” position. Subsequently, the donor substrate UDP-L-Ara4FN binds into the same GT-A domain, and UndP moves to position 2 (P2), the “catalysis” position. The nucleophilic attack is rapidly accomplished and soluble UDP is released, while the new UndP-L-Ara4FN is released back into the inner bacterial membrane. **B**) The proposed ArnC reaction mechanism involves a direct nucleophilic attack from the acceptor phosphate to the C1 of the L-Ara4FN ring. D100 acts as the catalytic base that abstracts a proton from the protonated oxygen of the UndP phosphate to enable the direct nucleophilic attack. The one step reaction enables the inversion of the anomeric configuration of the sugar. Several other residues that participate in coordination of the substrates are shown (D48, N77, D102, Q104, R128, R131, R137, R205).

Finally, we propose an overall mechanistic scheme for the catalytic cycle of ArnC in **Fig. 7A**. In this scheme, the acceptor UndP threads through the JM helices of an ArnC protomer, and is coordinated in position 1 (P1), the “standby” position. Subsequently, a donor substrate UDP-L-Ara4FN binds to the same GT-A and triggers a conformational rearrangement in the flexible β7-JM2 loop, which allows UndP to move to position 2 (P2), the “catalysis position” for the reaction shown in Fig. 7B to take place. After that, the newly formed product will force UndP to backtrack into a “product position”, which will facilitate release of the lipid back to the membrane. This is based on the observations that: i) UndP is coordinated stably within the GT-A domain in position 1 (**Fig. 5C**, **6A** and **S14B**) in the absence of a UDP nucleotide; ii) In the presence of the nucleotide both UndP and UDPA are less labile when UndP is located in position 2 (**Fig. 6B,C** and **S14C**); iii) Position 2 is optimal for catalysis to take place, as the acceptor phosphate is positioned in relative proximity of both the potential catalytic base D100 and the anomeric carbon of the L-Ara4FN sugar (**Fig. S14C,D**); iv) The “product position” for *Pf*DPMS shown in Fig. S14A is different that position 1, and since the DolP-Man structure (PDM 5MM1) is a product-bound state, it likely represents a temporary coordinating position of the product. The unbinding of the UDP nucleotide after catalysis will likely trigger the product to be expelled back into the membrane via the JM helices. As a final note, there is no indication that the binding of UndP in position 1 always happens first, in fact the binding of the donor and the acceptor is likely independent. Yet, the threading of UndP via the JM helices is expected to be a slower process compared to binding of a diffusible ligand. Thus, in most cases the binding of UndP will happen first and be rapidly succeeded by binding of the donor substrate and catalysis.

## Discussion

In this study we report the successful structure determination of the polyprenyl phosphate glycosyltransferase ArnC from the Gram-negative bacterium *S. enterica* embedded in lipid nanodiscs, in two conformations (*apo* and UDP-bound), using single-particle cryo-EM. We use MST to show that Mn^2+^ enables higher affinity for the partial donor substrate UDP, and by comparing the two conformations we show that binding of UDP on ArnC triggers a conformational rearrangement leading to a clamshell-like motion that brings the GT-A domain closer to the juxtamembrane helices of each protomer. We also perform a side-by-side comparison of two *apo* ArnC datasets collected using duplicate grids on a 200kV Talos Arctica and a 300kV Titan Krios microscope. At matched particle counts, datasets collected at 200 kV and 300 kV differed negligibly in resolution. No resolution cutoff was detected, as particle/resolution scaling conformed to theory throughout the range, but specimen-dependent limitations impose practical limits to resolution via the B-factor. Possible ways to improve the resolution of ArnC include alternative reconstruction algorithms, stabilizing or enlarging the complex, or physical improvements to the prepared grid. Next, we used CG simulations to show that the acceptor lipid UndP threads between the juxtamembrane helices of each ArnC protomer to reach the catalytic GT-A domain. Using additional atomistic simulations of ArnC with both the acceptor UndP and the full-donor substrate UDP-L-Ara4FN, we describe two different coordination positions for UndP within the GT-A domain (P1 and P2). We propose that the first position works as a “standby” position and the second as the “catalysis” position that enables the nucleophilic attack from the acceptor phosphate to the C1 carbon of the sugar. We also propose that the first aspartate of the **D**XD motif, which doesn’t participate in coordination of the metal ion, functions as a catalytic base to abstract a proton from the UndP and activate it to perform the nucleophilic attack. The proposed catalytic mechanism likely operates similarly on all members of the Pren-P GT family to enable partially deprotonated UndP molecules to react with the donor substrate.

Overall, the findings of this study provide extensive characterization of the structure, conformational changes, and mechanistic basis of function for the Pren-P GT ArnC, the third member of the family to be structurally elucidated after GtrB and *Pf*DPMS. Moreover, the structures of ArnC add to the repertoire of structurally characterized proteins in the aminoarabinose biosynthetic pathway that can be targeted for rational drug design aimed at preventing or reversing resistance to polymyxin antibiotics.

## Methods

### Target cloning

*arnC* genes from the clinically relevant species *S. enterica* (WP_000458893), *K. pneumoniae* (WP_040165138), and *E. coli* (NP_416757) were codon optimized for expression in *E. coli* and synthesized with 30bp overhangs (Genewiz) for direct assembly into expression vectors using NEBuilder^®^ HiFi DNA Assembly Master Mix (NEB E2621). The genes were cloned in frame with either an N-terminal FLAG-10xHis-TEV cassette or a C-terminal TEV-10xHis cassette using pNYCOMPS vectors^43^. The sequences of the synthetic linear DNA fragments and the primers used to amplify the two pNYCOMPS vectors (N-term, C-term) with PCR are shown in **Table 2**.

**Table 2.**
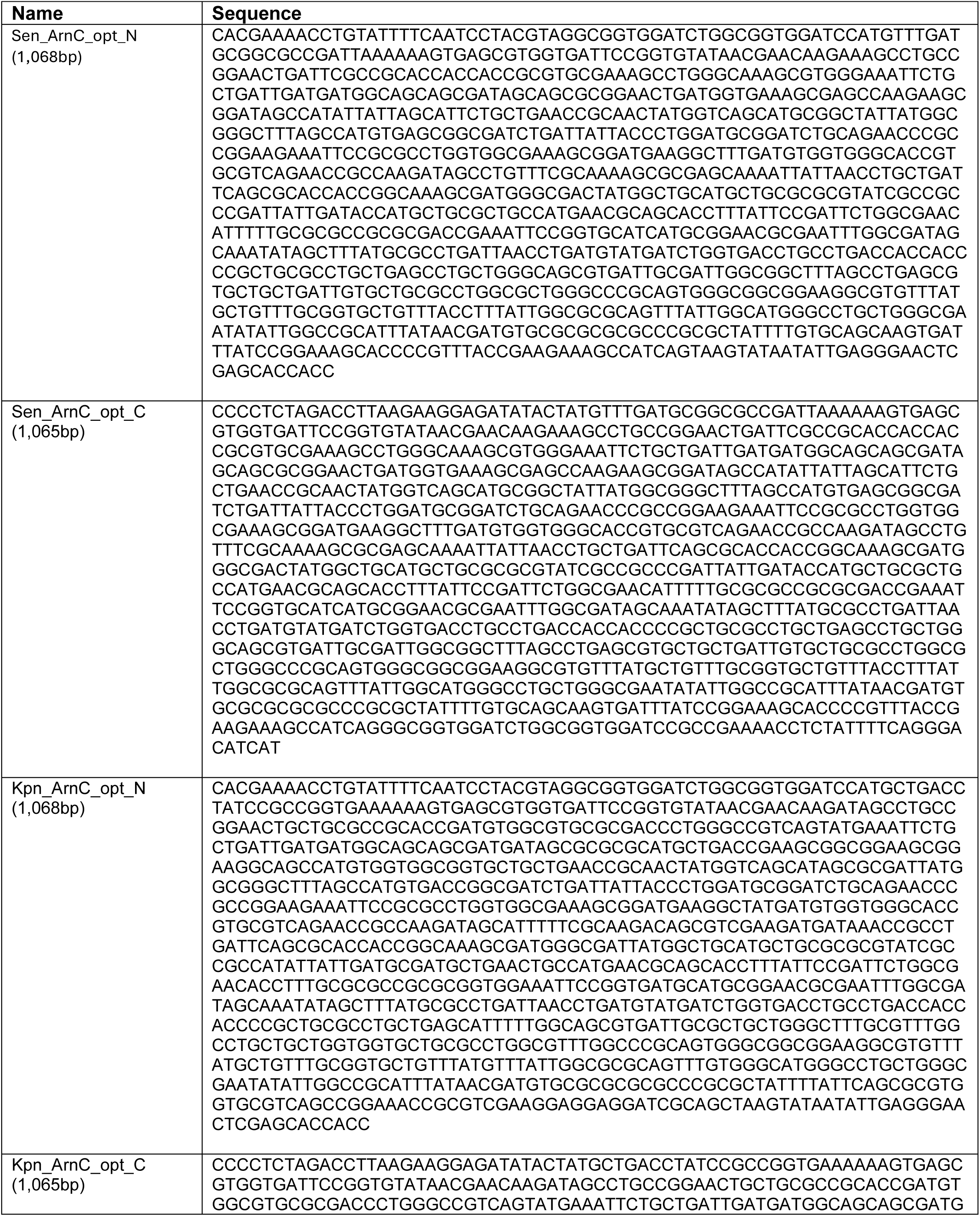
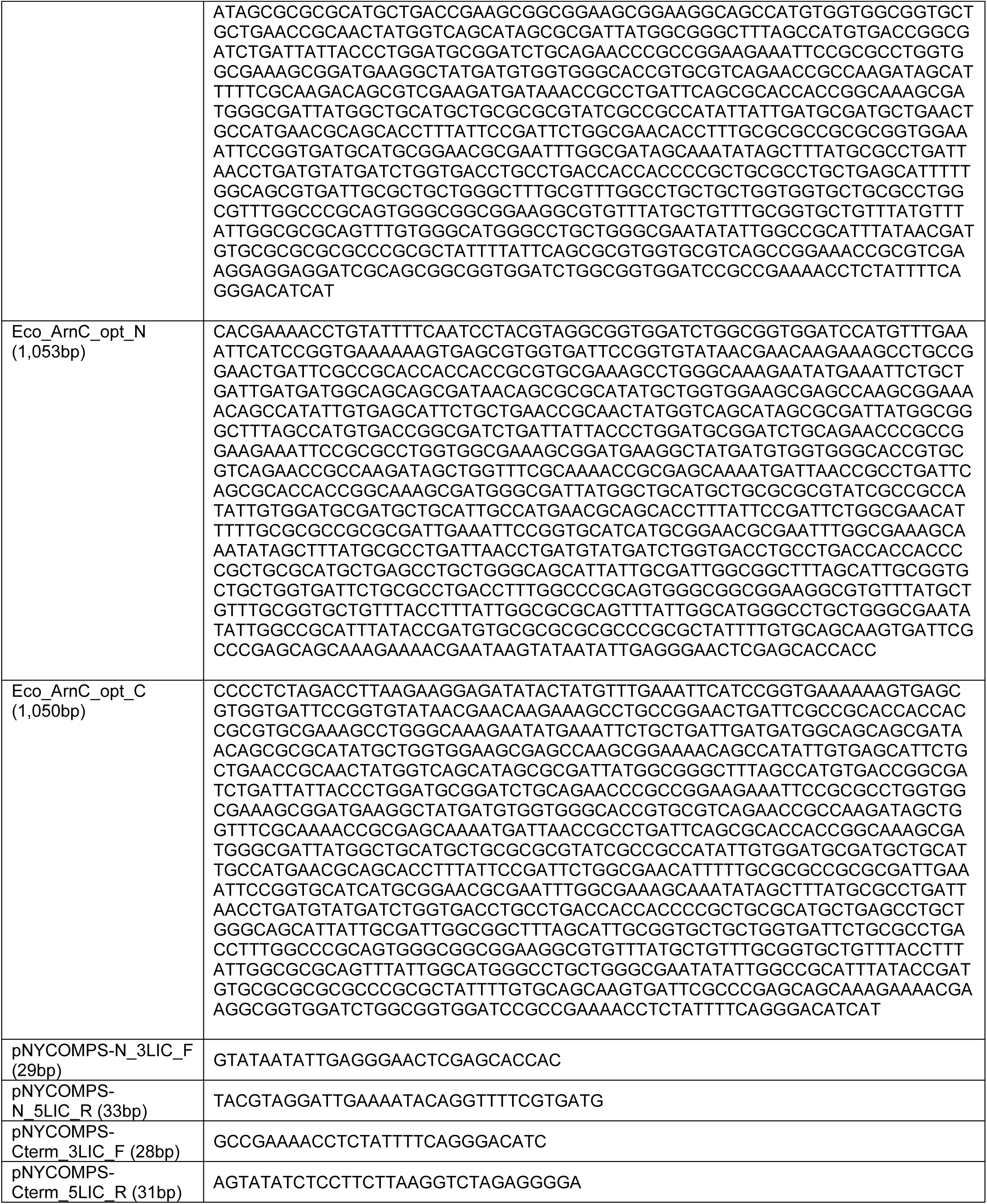
Linear DNA fragments and primers used for *arnC* gene cloning.

### Small-scale expression and purification

The resulting ArnC constructs were transformed into T7 Express lysY E. coli competent cells (NEB, C3010I), an enhanced BL21 derivative strain, and grown overnight in 1mL 2xYT medium supplemented with 50μg/mL kanamycin at 37°C with shaking (240 RPM). The next day, 20mL of the same medium was inoculated with the starter culture at 1:100 ratio and grown at 37°C with shaking (240 RPM), until OD600 1.0 was reached. Temperature was then decreased to 22°C and protein expression was induced with 0.2mM isopropyl β-D-1-thiogalactopyranoside (IPTG), and the culture was incubated overnight with shaking (240 RPM). Cells were harvested by centrifugation (4,000g for 10 min) at 4°C. For small scale purification of ArnC proteins, the cell pellets were resuspended in 700µL lysis buffer containing 20mM HEPES pH 7.5, 200mM NaCl, 20mM MgSO4, 10µg/mL DNase I, 8µg/mL RNase A, 1 mM tris(2-carboxyethyl)phosphine hydrochloride (TCEP), 1mM PMSF, and protease inhibitor cocktail set III EDTA-free (Calbiochem, 539134) at 1:1000 ratio. Cells were lysed using a probe sonicator (Microson) and the lysate was solubilized for 1.5 hour with n-dodecyl-β-D-maltopyranoside (DDM; Anatrace, D310S) at a final concentration of 1% (w/v). Insoluble materials were removed by centrifugation at 18,000g for 30 min at 4°C. The supernatant was then subjected to metal-affinity chromatography using NiNTA agarose beads (Qiagen, 30230). Briefly, the supernatants were incubated with pre-equilibrated NiNTA agarose beads overnight (25µL per pellet from a 20mL culture). The beads were loaded on a 96-well filter plate and washed thrice with 100µL of wash buffer containing 20mM HEPES pH 7.5, 500mM NaCl, 80mM imidazole and 0.1% (w/v) DDM. The flow through at each step was removed by brief centrifugation (100g for 30 sec) at 4°C. ArnC proteins were eluted with 60µL elution buffer containing 20mM HEPES pH 7.5, 200mM NaCl, 300mM imidazole, and 0.1% (w/v) DDM. the eluted proteins were run on 12% SDS-PAGE gel and stained with Imperial protein stain (Thermo Scientific 24617) for visualization (**Fig. S1A**).

### Large-scale expression and purification in detergent

For large scale production of ArnC from Salmonella enterica (ArnC*_Se_*), the construct encoding N-terminal FLAG-10xHis-TEV-ArnC*_Se_* was used to transform T7 Express lysY E. coli competent cells (NEB, C3010I), and grown overnight in 2xYT medium supplemented with 50μg/mL kanamycin at 37°C with shaking (240 RPM). The next day, 200mL of the same medium were inoculated with the starter culture at 1:100 ratio and left to grow at 37°C with shaking (240 RPM), until OD600 reached 0.8-1.2. Temperature was then reduced to 22^°^C, protein expression was induced with 0.2mM IPTG, and the culture was incubated overnight with shaking (240 RPM). Cells were harvested by centrifugation (3,700 RPM for 15 min) at 4°C, washed once with phosphate buffered saline (PBS) and centrifuged again to produce a solid pellet that was stored at –80°C until further use. For large-scale purification cell pellets were resuspended in lysis buffer containing 20mM HEPES pH 7.5, 200mM NaCl, 20mM MgSO4, 10μg/mL Dnase I, 8μg/mL Rnase A, 1mM TCEP, 1mM PMSF, and protease inhibitor cocktail set III EDTA-free (Calbiochem, 539134) at 1:1000 ratio. Cells were lysed with an Emulsiflex C3 high-pressure homogenizer (Avestin) and the lysate was solubilized for 2h with DDM (Anatrace, D310S) added to a final concentration of 1% (w/v), in a volume of approximately 40mL per cell pellet from 800mL culture. Insoluble material was removed by ultracentrifugation at 34,000 RPM for 30 min at 4°C and the protein was purified from the supernatant by metal-affinity chromatography using Ni-NTA agarose beads (Qiagen, 30230). The supernatant, after addition of 40mM imidazole, was incubated with pre-equilibrated Ni-NTA agarose beads (0.7mL per pellet from an 800mL culture) overnight. The beads were then loaded on a column and washed with 10 column volumes of 20mM HEPES pH 7.5, 500mM NaCl, 75mM imidazole and 0.03% (w/v) DDM. Protein was eluted with 4 column volumes of 20mM HEPES pH 7.0, 150mM NaCl, 300mM Imidazole, and 0.03% (w/v) DDM. Imidazole was removed from the eluted protein by exchanging buffer to 20mM HEPES pH 7.0, 200mM NaCl, 0.03% (w/v) DDM (final protein buffer) using a PD-10 desalting column (Cytiva, 17085101).

### Nanodisc incorporation

ArnC was incorporated into lipid nanodiscs with a 1:300:5 molar ratio of protein:1-palmitoyl-2-oleoyl-sn-glycero-3-phospho-(1’-rac-glycerol) (POPG):membrane scaffold protein 1E3D1 (MSP1E3D1)^59,60^. This mixture was incubated at 4°C for 2h with gentle agitation. Reconstitution was initiated by removing detergent with the addition of Bio-beads SM-2 (Bio-Rad, 1523920) at 4°C overnight with constant rotation. Subsequently, bio-beads were removed and the nanodisc reconstitution mixture was bound again to Ni-NTA resin at 4°C for 2h to remove empty nanodiscs. The resin was washed with 10 column volumes of wash buffer (20mM HEPES pH 7.5, 150mM NaCl, and 60mM imidazole) followed by four column volumes of elution buffer (20mM HEPES pH 7.0, 150mM NaCl, and 300mM imidazole). The sample was further purified by loading onto a Superdex 200 Increase 10/300 GL size-exclusion column (Cytiva, 28990944) in gel filtration buffer (20mM HEPES pH 7.0 and 150mM NaCl). Protein typically eluted as a sharp monodispersed peak, observed by monitoring absorbance at 280 nm (**Fig. S1C**).

### Microscale Thermophoresis (MST)

Prior to labeling, detergent-purified ArnC was diluted to 20µM in buffer A (20mM HEPES, pH 7.5, 150mM NaCl, 0.01% DDM). The diluted ArnC was labeled using a Monolith Protein Labeling Kit RED-NHS 2nd Generation (Amine reactive) (NanoTemper Technologies, MO-L011). To assess the binding affinity of ArnC for UDP in the presence of different metal ions, UDP was titrated into ArnC across four experimental conditions: 1mM MnCl₂, 1mM MgCl₂, 1mM EDTA to chelate any endogenous metal ions, or without adding any exogenous metal ions. ArnC was used at a final concentration of 50nM. The UDP titration was prepared by serial dilution in each condition, starting from 2mM to 122nM. After incubation, the samples were transferred into premium coated capillaries (Nanotemper Technologies, MO-K025) and read in a Monolith NT.115 Nano-Blue/Red instrument at room temperature using 40% light-emitting-diode (LED) power and 40% MST power. Binding affinities were calculated using the thermophoresis with T jump evaluation strategy. All experiments were repeated three times in duplicate (n=6) for each experimental group. Data analyses were performed using the NanoTemper analysis software.

### Single-particle cryo-EM vitrification and data acquisition

Purified ArnC incorporated into MSP1E3D1/POPG nanodiscs was concentrated to 1.5mg/ml using an Amicon 100kDa concentrator (EMD Milipore, UFC510096). To generate UDP-bound ArnC, the purified protein was laced with 2mM Uridine 5-Diphosphate Disodium (UDP) (Sigma-Aldrich, 94330) and 1mM MnCl_2_, prior to freezing. For each sample, 3μl of sample was added to a glow-discharged R 1.2/1.3, 300 mesh holey gold grid (Quantifoil, UltrAuFoil) and blotted using filter paper for 3.5 s using a Vitrobot Mark IV (ThermoFisher Scientific) with a blot force of 3 and a wait time of 30 sec, before plunging into liquid ethane for vitrification. The sample chamber was held at 4°C with >90% humidity to minimize evaporation and sample degradation.

### *apo* ArnC data collection on Talos Arctica

Micrograph movies in TIFF format were collected at the Rutgers Cryo-EM & Nanoimaging Facility (RCNF) using a 200 kV Talos Arctica electron microscope (FEI/ThermoFisher Scientific), equipped with a K2 Summit direct electron detector (Gatan) and a BioQuantum energy filter (Gatan) with slit width of 20 eV. Data were collected automatically in counting mode using EPU (FEI/ThermoFisher Scientific), a nominal magnification of 130,000×, a nominal pixel size of 1.038 Å/pixel, and a dose rate of 5.420 electrons/pixel/s. Movies were recorded at 200 ms/frame for 7s (35 frames total), resulting in a total dose of 35.21 electrons/Å^2^. Nominal defocus range was −0.5 to −2.5 μm. A total of 5,552 movies were recorded from one grid over three days. Micrographs were gain-normalized and defect-corrected. At the end of processing a calibrated pixel size of 1.05 Å/pixel was determined for this dataset, representing an 1.16% error from the nominal pixel size value. The dataset has been deposited to EMPIAR with accession code EMPIAR-11924.

### *apo* ArnC data collection on Titan Krios

Micrograph movies in TIFF format were collected at the National Center for Cryo-EM Access and Training (NCCAT) using the 300kV Titan Krios #4 – “Elizabeth” electron microscope (ThermoFisher Scientific), equipped with a K3 direct electron detector (Gatan) and a BioQuantum energy filter (Gatan) with slit width of 20 eV. Data were collected automatically in counting mode using image shift in Leginon, a nominal magnification of 130,000×, a nominal pixel size of 0.646 Å/pixel, and a dose rate of 19.9685 electrons/pixel/s. Movies were recorded at 30 ms/frame for 1.2s (40 frames total), resulting in a total dose of 57.42 electrons/Å^2^. Nominal defocus range was −0.8 to −2.0 μm. A total of 23,259 movies were recorded from one grid over three days. Micrographs were not gain-normalized or defect corrected. A gain reference was recorded separately and is provided together with the dataset. At the end of processing a calibrated pixel size of 0.67 Å/pixel was determined for this dataset, representing an error of 3.72% from the nominal pixel size value. The dataset has been deposited to EMPIAR with accession code EMPIAR-12002.

### UDP-bound ArnC data collection on Talos Arctica

Micrographs in TIFF format were collected at RCNF using a 200 kV Talos Arctica electron microscope (FEI/ThermoFisher Scientific), equipped with a K2 Summit direct electron detector (Gatan) and a BioQuantum energy filter (Gatan) with slit width of 20 eV. Data were collected automatically in counting mode using EPU (FEI/ThermoFisher Scientific), a nominal magnification of 165,000×, a nominal pixel size of 0.818 Å/pixel, and a dose rate of 5.0344 electrons/pixel/s. Movies were recorded at 100 ms/frame for 5.3s (53 frames total), resulting in a total dose of 39.88 electrons/Å^2^. The nominal defocus range used was −0.5 to −2.5 μm. A total of 7,925 micrographs were recorded from one grid over two days. Micrographs were gain-normalized and defect corrected. At the end of processing a calibrated pixel size of 0.82 Å/pixel was determined for this dataset, representing an 0.24% error from the nominal pixel size value. The dataset has been deposited to EMPIAR with accession code EMPIAR-11987.

### Cryo-EM data processing

For the Arctica *apo* ArnC dataset: Motion correction and contrast transfer function (CTF) estimation were performed using patch motion correction and patch CTF estimation as implemented in cryoSPARC (v.3.3.2)^61^. The resulting aligned micrographs were manually curated for CTF fit resolution (<6Å), defocus (>5500Å), average intensity (−35<intensity<150), relative ice thickness (<3), and total full-frame motion distance (<45pix), resulting in 3,402 exposures (∼61%) being accepted. Blob picker (min diameter 60Å, max diameter 150Å) was used on 500 random micrographs to pick particles, and 50,752 particles were extracted using a 256-pixel box size and binned 4X. The particle stack was then subjected to 2D classification in cryoSPARC v.3.3.2. Eight 2D classes corresponding to 8.585 particles where selected and used for template picking of 600 random micrographs to pick particles.

78,806 particles were extracted using a 288-pixel box size binned 4X. The particle stack was again subjected to 2D classification and seven 2D classes corresponding to 24,123 particles were used for template picking the full 3,402 accepted micrograph dataset. A total **1,002,867** particles were extracted using a 288 pixel box with 3X binning. This particle stack was subjected to 3 rounds of cleanup using *ab initio* reconstruction with 3 classes, resulting in a particle stack of 224,496 that was re-extracted with 288 pixel box size with no binning. This particle stack was subjected to a series of *ab initio* and heterogeneous refinement steps, resulting in a 3.42 Å map, which was then subjected to non-uniform refinement and local refinement in cryoSPARC resulting in a 2.73 Å reconstruction map from a final particle stack of 187,649 particles after imposing C4 symmetry. A calibrated pixel size of 1.05 Å/pix was imposed on the final reconstruction giving a final resolution of 2.79Å. The processing workflow is summarized on **Fig. S2A**.

For the Krios *apo* ArnC dataset: Motion correction and CTF estimation was performed using patch motion correction and patch CTF as implemented in cryoSPARC (v.3.3.2). The micrographs were separated to 4 groups of ∼4-5K micrographs each. The resulting aligned micrographs were manually curated for CTF fit resolution (<5.9Å), defocus (>5500Å), average intensity (−7<intensity<4.4), relative ice thickness (<1.5), and total full-frame motion distance (<60pix), resulting in 17,547 exposures (∼75%) being accepted. Templates were imported from the *apo* Arctica dataset and used for template picking of each group of 4K micrographs. Approx. 1m particles per group was extracted using a 384-pixel box size binned 4X, which were then subjected to 2 separate rounds of *ab initio* cleanup, The resulting particle stack containing 525,744 particles was re-extracted using a 480-pixel box size (no binning). Heterogeneous refinement with 1 model and 3 decoys gave a particle stack of 506,322 particles. This particle stack was subjected to non-uniform refinement, manual micrograph curation, and then local refinement in cryoSPARC resulting in a 2.62 Å reconstruction map from a final particle stack of 490,807 particles after imposing C4 symmetry. A calibrated pixel size of 0.67 Å/pix was imposed on the final reconstruction giving a final resolution of 2.74Å. The processing workflow is summarized on **Fig. S3A**.

For the UDP-bound ArnC dataset: Motion correction and CTF estimation was performed using patch motion correction and patch CTF as implemented in cryoSPARC (v.3.3.2). The resulting aligned micrographs were manually curated for CTF fit resolution (<6.8Å), defocus (>1400Å), average intensity (−45<intensity<100), astigmatism<1000, relative ice thickness (<1.26), and total full-frame motion distance (<35pix), resulting in 6,761 exposures (∼85%) being accepted. Templates were imported from the *apo* Arctica dataset and used for template picking. 2,382,682 particles were extracted using a 288-pixel box size binned 4X. This particle stack was then subjected to separate rounds of *ab initio* and 2D classification cleanup resulting in a particle stack of 347,008 particles that were reextracted using an unbinned 288-pixel box size. This particle stack was again subjected to a series of *ab initio* reconstructions with 2 classes, heterogeneous and homogeneous refinement, and manual micrograph curation, resulting in a 2.87Å map from 216,104 particles after imposing C4 symmetry. A calibrated pixel size of 0.82 Å/pix was imposed on the final reconstruction giving a final resolution of 2.96Å. The processing workflow is summarized on **Fig. S9A**.

### Pixel calibration

The calibrated pixel size for each dataset was determined using aa 1-120 of the GtrB X-ray crystal structure (PDB 5EKP) to fit the density of the GT-A domain in ArnC. Voxel size was varied systematically using UCSF Chimera and the correlation was monitored until it was maximized. The voxel size at the maximal correlation was selected as the calibrated pixel size and was imposed on the final reconstruction for each dataset.

### Atomic model building and refinement

For modeling, an AlphaFold generated model of the E. coli (K12 strain) ArnC (Uniprot: P77757) from the AlphaFold database^47^ was fitted in the *apo* (Arctica) 2.79 Å cryo-EM map using UCSF Chimera, and was manually mutated in Coot^62^ to match the sequence of *S. enterica* ArnC. Subsequent model refinement and adjustment was performed in Coot^62^, Phenix^63^ and Isolde^64^, iteratively. The *apo* (Arctica) model was then used to build models for the *apo* (Krios) map and UDP-bound cryo-EM maps. UDP was manually placed into the observed density using Coot^62^ and refined in Phenix^63^. After refinement of each model in Isolde^64^, a final refinement using phenix.real_space_refine was performed before finalizing each structure. The program phenix.douse was used to add waters automatically to the *apo* ArnC structure from the Krios map. Waters were manually curated in a second step before a final refinement using phenix.real_space_refine.

### Resolution analysis

Random subsets of particles were reconstructed in cryoSPARC (v.3.3.2.). Orientational assignments, defoci, and other particle parameters from the final 3D refinement job (*vide supra*) were preserved; no refinement was performed. Only the 3D reconstruction step was repeated using the job type “Homogeneous Reconstruction Only.” Spectral signal-to-noise ratio was calculated as the Fourier shell correlation between the even and odd half-maps. Radial averaging of structure factors was performed using EMAN^65^, and simulated maps were formed by approximating each atomic position as a sphere of density with Gaussian spatial decay.

### Model analysis

A cavity search using the Solvent Extractor from Voss Volume Voxelator server^66^ was performed using an outer-probe radius of 10 Å and inner-probe radius of 2 Å. Chimera^67^ and ChimeraX^68^ were used to visualize the structures and the resulting cavity volumes.

### Coarse-grained MD simulations

Coarse-grained MD simulations of ArnC were established using the MemProtMD^69,70^ Google Colab Notebook and LipIDens^55^ for the apo ArnC conformation, including the modelling of missing residues/loops. ArnC was embedded in a bilayer composed of POPE (64%), POPG (18%), cardiolipin (CDL) (9%) and UndP (9%) and solvated with MARTINI water^71^ and 0.15 M NaCl in a 12 x 12 x 12 nm3 box. The Martini 3.0 forcefield was used to describe all components^71^. Systems were energy minimized using a steepest-descent method and equilibrated in 2 x 1 ns steps. 10 x 15 μs CG simulations were run using parameters from LipIDens^55^ and the GROMACS 2020 simulation software^72^. Temperature was maintained at 310 K using the V-rescale thermostat^73^ (*τ*t= 1.0 ps) and pressure was maintained at 1 bar using the Parrinello-Rahman barostat^74^ (*τ*p= 12.0 ps, compressibility = 3 x 10-4). The timestep was 20 fs. Coulombic interactions were described using the reaction-field method and a 1.1 nm cut-off. Van der Walls interactions were described with the potential-shift Verlet method and a 1.1 nm cut-off. Lipid interactions were analysed using the PyLipID toolkit^75^ within the LipIDens pipeline^55^. A range of dual cut-off schemes were sampled and a 0.5/0.7 nm dual cut-off was selected based on the lipid interaction probability distribution and screening of e.g. interaction durations and binding site numbers across cut-off schemes. Lipid interaction residence times and binding sites were calculated using PyLipID^75^. Site screening and comparisons were obtained using LipIDens with POPE as the reference lipid against which to compare sites.

### Atomistic MD simulations

The top ranked binding pose for UndP bound within the ArnC GT-A domain was selected for simulation. The LipIDens pipeline employed the CG2AT2 tool^76^ to backmap the CG frame to atomistic resolution. The protein conformation was backmapped to that within the apo structure. The CHARMM-36 forcefield was used to describe all components^77^. Water was described using the TIP3P model^78^. Atomistic simulations were run for 3×100 ns using GROMACS 2020^72^ and parameters from LipIDens^55^. Temperature was maintained at 310 K using the Nosé-Hoover thermostat^79^ (*τ*t= 0.5 ns) and pressure was maintained at 1 bar using the Parrinello-Rahman barostat^74^ (*τ*p= 2.0 ps, compressibility = 4.5 x 10-5). Van der Walls interactions were switched from 1.0 nm to 1.2 nm with the force-switch modifier. Coulombic interactions were modelled with the Particle-Mesh Ewald method^80^ and a 1.2 nm cut-off. The LINCS algorithm^81^ was used to constrain bonds to their equilibrium values and a dispersion correction was not applied. Per residue interaction occupancies with the bound UndP were calculated using a 0.3 nm cut-off. PyMol, ChimeraX and VMD were used for visualisation^68,82,83^.

### Molecular modelling of substrate-bound ArnC and atomistic simulations

The UDP-bound E. coli ArnC structure was used to build molecular models for the interactions with substrates, UndP and UDP-L-Ara4FN (UDPA), and Mg2+. While the coordinates for UDP portion of UDPA were modelled as in the cryo-EM structure, two conformations for the Ara4FN group of UDP-L-Ara4FN and the UndP were proposed (Figure 6A). The first model is based on the coordination of the phosphate of UndP by the conserved arginines R128 and R137 as suggested by the Coarse-Grained simulations. For this model, the Ara4FN sugar was placed in an open groove in a distance from the phosphate of UndP that would allow the reaction to proceed (Figure 6A – State 1). A second model was generated using the same coordinates for UndP as in State 1, while the position of the Ara4FN group was based on the structure of *Pf*DPMS in complex with GDP-mannose (PDB ID: 5MM0)^42^ (Figure 6A – State 2). For this model, UndP and UDPA are positioned in a distance that would not allow the reaction to occur; therefore a third model was proposed in which UDPA is positioned as in the previous model and lipid is placed further down in the active site with the phosphate group of UndP below the sugar (Figure 6A – State 2).

Atomistic simulations of substrates bound to ArnC were performed using the CHARMM36m force field^77^. As parameters for Mn^2+^ are not readily available for the CHARMM36m force field, so Mg^2+^ was used for the atomistic simulations. Parameters for UndP and UDP were taken from the CHARMM36m force field. CHARMM-GUI was used to prepare the CHARMM36m parameters and coordinates for UndP-Ara4FN and UDP-Ara4FN.

The MemProtMD pipeline^69,70^ was used to run an initial 1 μs CG MD simulation to permit the assembly and equilibration of a POPG:POPE bilayer around ArnC without its substrate. The input protein was aligned to *xy* plane using MEMEMBED^84^ and then converted to a CG representation using the Martini 3 force field^71^. In brief, the ArnC model was placed at the center of a periodic box and a POPG-POPE bilayer at 1:4 molar ratio was built around the protein. The final snapshot was converted back to atomic details using CG2AT^76^ and the substrates coordinates, as described above, were added to the system. NaCl was added in a concentration of 150 mM to render the systems neutral. The complete systems were further equilibrated for 1 ns maintaining the structure of the protein and ligands restrained. Three repeats of unrestrained 500 ns MD simulations were performed for each system. All simulations were performed in the isothermal-isobaric ensemble at 310 K and 1 bar using a timestep of 2 fs. Pressure was maintained at 1 bar with a semi-isotropic compressibility of 4 x 10-5 using the Parrinello-Rahman barostat^74^. Temperature was controlled using the velocity-rescale thermostat^73^, with the solvent, lipids and protein coupled to an external bath. The long-range electrostatic interactions were computed with the Particle Mesh Ewald method^80^, while a verlet cut-off method was used to compute the non-bonded interactions. All MD simulations were performed using GROMACS^72^ 2022 and analyzed using GROMACS tools and MDAnalysis^85^. All images were generated using PyMOL^82^.

## Data Availability

Atomic coordinates of the cryo-EM structures generated in this study have been deposited to the Protein Data Bank (PDB) under accession numbers: 8VXH (*apo* – Arctica), 9B77 (*apo* – Krios), and 9ASC (UDP – Arctica). Cryo-EM maps corresponding to the final sharpened map and half-maps for each dataset have been deposited to the Electron Microscopy Data Bank (EMDB) under accession numbers: EMD-43617 (*apo* – Arctica), EMD-44302 (*apo* – Krios), and EMD-43812 (UDP – Arctica). Raw electron micrograph movies, aligned and dose-weighted micrographs, and the final particle stacks used in reconstructions have been deposited to the Electron Microscopy Public Image Archive (EMPIAR) under accession numbers: EMPIAR-11924 (*apo* – Arctica), EMPIAR-12002 (*apo* – Krios), and EMPIAR-11987 (UDP – Arctica). Additional atomic models used in this study are publicly available under accession codes 5EKP, 5MLZ, 5MM0, 5MM1.

## Acknowledgements

This project was supported by NIH grants R00GM123228 and R35GM150831 to VIP. PJS acknowledges the NIH (R01AI174416 - PI: M. Stephen Trent), Wellcome (208361/Z/17/Z), MRC, BBSRC, and the Howard Dalton Centre for funding. PJS and CMB acknowledge Sulis at HPC Midlands+, which was funded by the EPSRC on grant EP/T022108/1, and the University of Warwick Scientific Computing Research Technology Platform for computational access. This project made use of time on ARCHER2 and JADE2 granted via the UK High-End Computing Consortium for Biomolecular Simulation, HECBioSim (http://www.hecbiosim.ac.uk), supported by EPSRC (grant no. EP/R029407/1). TBA was supported by Wellcome (102164/Z/13/Z) and is currently supported by Schmidt Science Fellows, in partnership with the Rhodes Trust. The Krios dataset was collected at the National Center for Cryo-EM Access and Training (NCCAT) and the Simons Electron Microscopy Center located at the New York Structural Biology Center, supported by the NIH Common Fund Transformative High Resolution Cryo-Electron Microscopy program (U24 GM129539) and by grants from the Simons Foundation (SF349247) and NY State Assembly. We thank the staff at NCCAT that facilitated the collection of the *apo* ArnC dataset.

---

**Figure S1.**
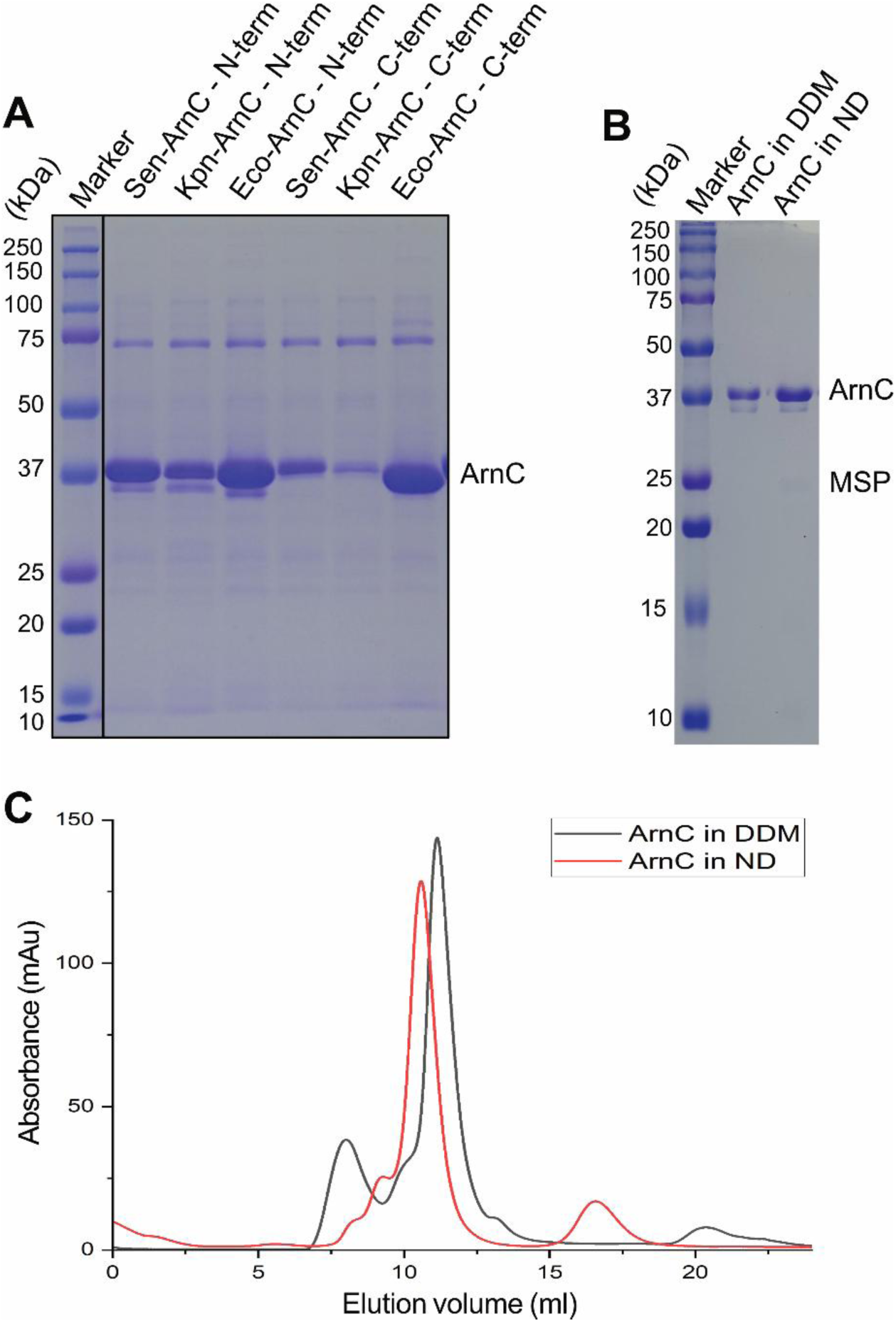
Expression and purification of *S. enterica* ArnC. **A**) SDS-PAGE gel showing relative expression of ArnCs from *Salmonella enterica* (*Sen*), *Klebsiella pneumoniae* (*Kpn*) and *Escherichia coli* (*Eco*), with either N-teminal or C-terminal expression tags. **B**) SDS-PAGE gel of ArnC from *S. enterica* large-scale purification, showing ArnC in DDM after affinity purification, and ArnC reconstituted into MSP1E3D1/POPG nanodiscs (ND). **C**) Size-exclusion chromatography elution profiles of purified ArnC in detergent (black), and incorporated into a nanodisc (red), on a Superdex 200 Increase 10/300 GL column (Cytiva).

**Figure S2.**
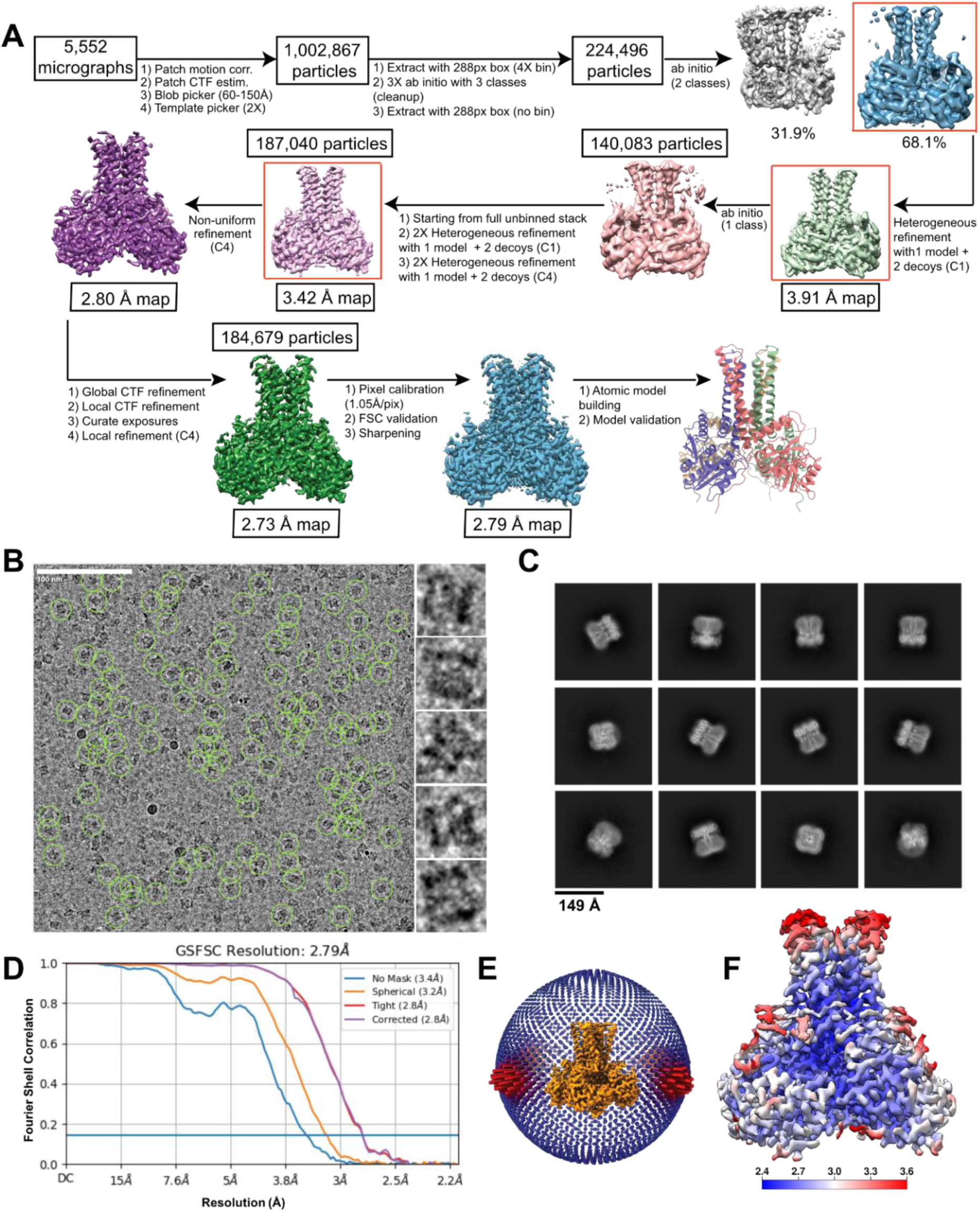
Cryo-EM analysis of apo ArnC*_Se_* collected on Talos Arctica. **A**) Data processing workflow used to determine the structure of nanodisc-reconstituted ArnC*_Se_* from a Talos Arctica dataset. **B**) Representative electron micrograph of nanodisc-reconstituted ArnC*_Se_* from Talos Arctica. Particles included in the final reconstruction are marked with green circles. Insets on the right show individual single particles from the micrograph. Scale bar, 100nm. **C**) Representative 2D class averages after reference-free 2D classification of the final particle stack in cryoSPARC. **D**) Fourier shell correlation (FSC) curve for apo nanodisc-reconstituted ArnC*_Se_* after the final local refinement in cryoSPARC. **E**) Euler angle distribution plot of all ArnC*_Se_* particles used in the final reconstruction. Final map shown in orange. Each orientation is represented by a cylinder, with each cylinder’s height and color (from blue to red) proportional to the number of particles for that specific orientation. **F**) Local resolution map for the final ArnC*_Se_* reconstruction. Local resolution was estimated using an implementation of the blocres program in cryoSPARC. Coloring shown from deep blue (2.4 Å) to red (≥3.6 Å).

**Figure S3.**
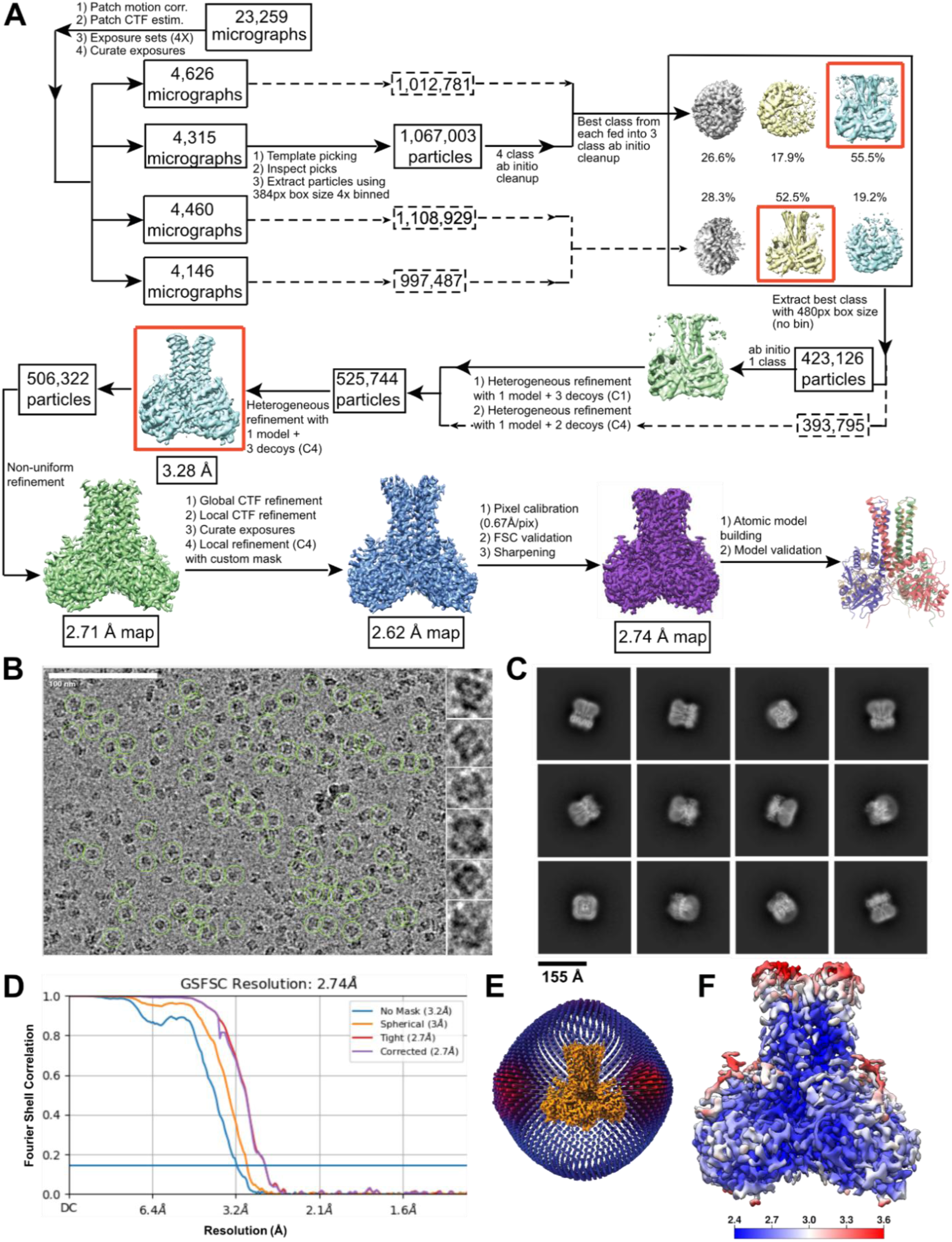
Cryo-EM analysis of apo ArnC*_Se_* collected on Titan Krios. **A**) Data processing workflow used to determine the structure of nanodisc-reconstituted ArnC*_Se_* from a Krios microscope dataset. **B**) Representative electron micrograph of ArnC*_Se_* from Titan Krios. Particles included in the final reconstruction are marked with green circles. Insets on the right show individual single particles from the micrograph. Scale bar, 100nm. **C**) Representative 2D class averages after reference-free 2D classification of the final particle stack in cryoSPARC. **D**) Fourier shell correlation (FSC) curve for apo nanodisc-reconstituted ArnC*_Se_* after the final local refinement in cryoSPARC. **E**) Euler angle distribution plot of all ArnC*_Se_* particles used in the final reconstruction. Final map shown in orange. **F**) Local resolution map for the final ArnC*_Se_* reconstruction. Coloring shown from deep blue (2.4 Å) to red (≥3.6 Å).

**Figure S4.**
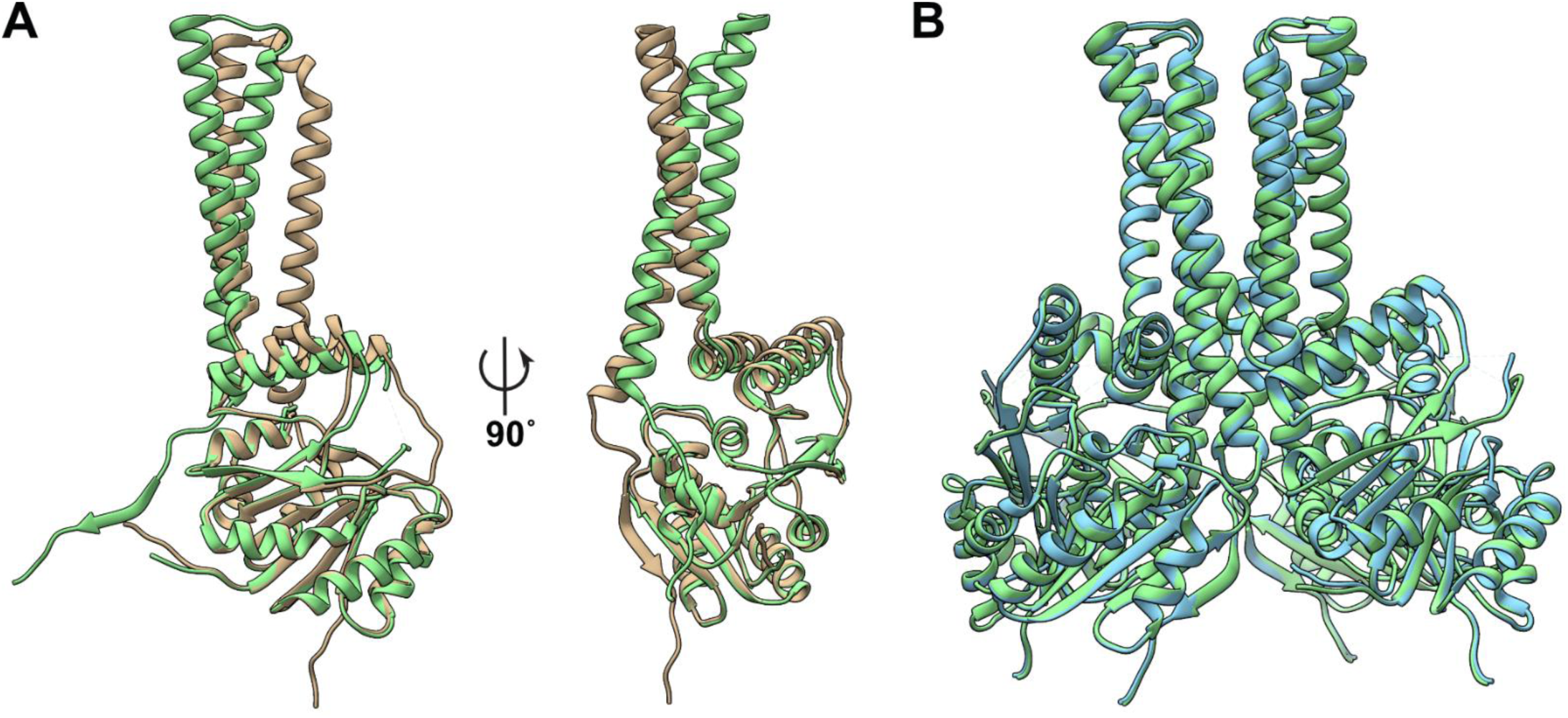
*apo* ArnC model comparison. **A**) Comparison between the final apo ArnC*_Se_* atomic model (green) and the Alphafold predicted atomic model of ArnC from *E. coli* (AF-P77757-F1-v4) used as a starting model for model building. RMSD between 312 atom pairs is 9.54Å. **B**) Comparison between the final apo ArnC_Se_ atomic model built based on the Arctica dataset map (green) and the one build based on the Krios dataset map (light blue). RMSD between 1248 atom pairs is 0.67Å.

**Figure S5.**
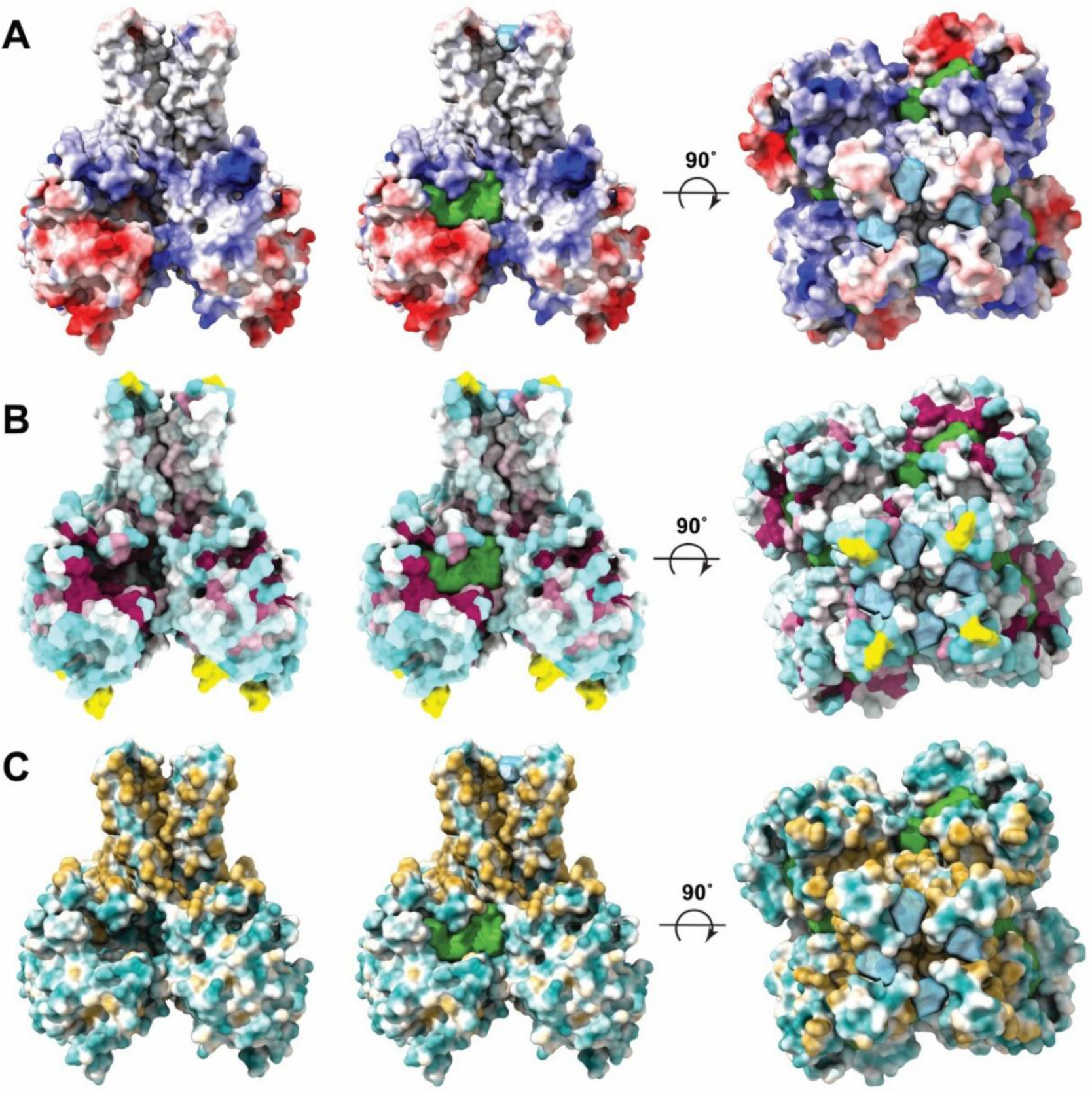
Physicochemical properties of ArnC. **A**) ArnC*_Se_* rendered in surface representation colored by electrostatic potential. **B**) ArnC*_Se_* surface colored by residue conservation on a yellow/white (no conservation) to purple (absolute conservation) scale. **C**) ArnC*_Se_* surface colored by Wimley-White hydrophobicity, on a cyan (very hydrophilic) to gold (very hydrophobic) scale. Two orthogonal views are presented with cavities shown either empty (left) or filled with a green (cavity 1) or light blue (cavity 2) volume.

**Figure S6.**
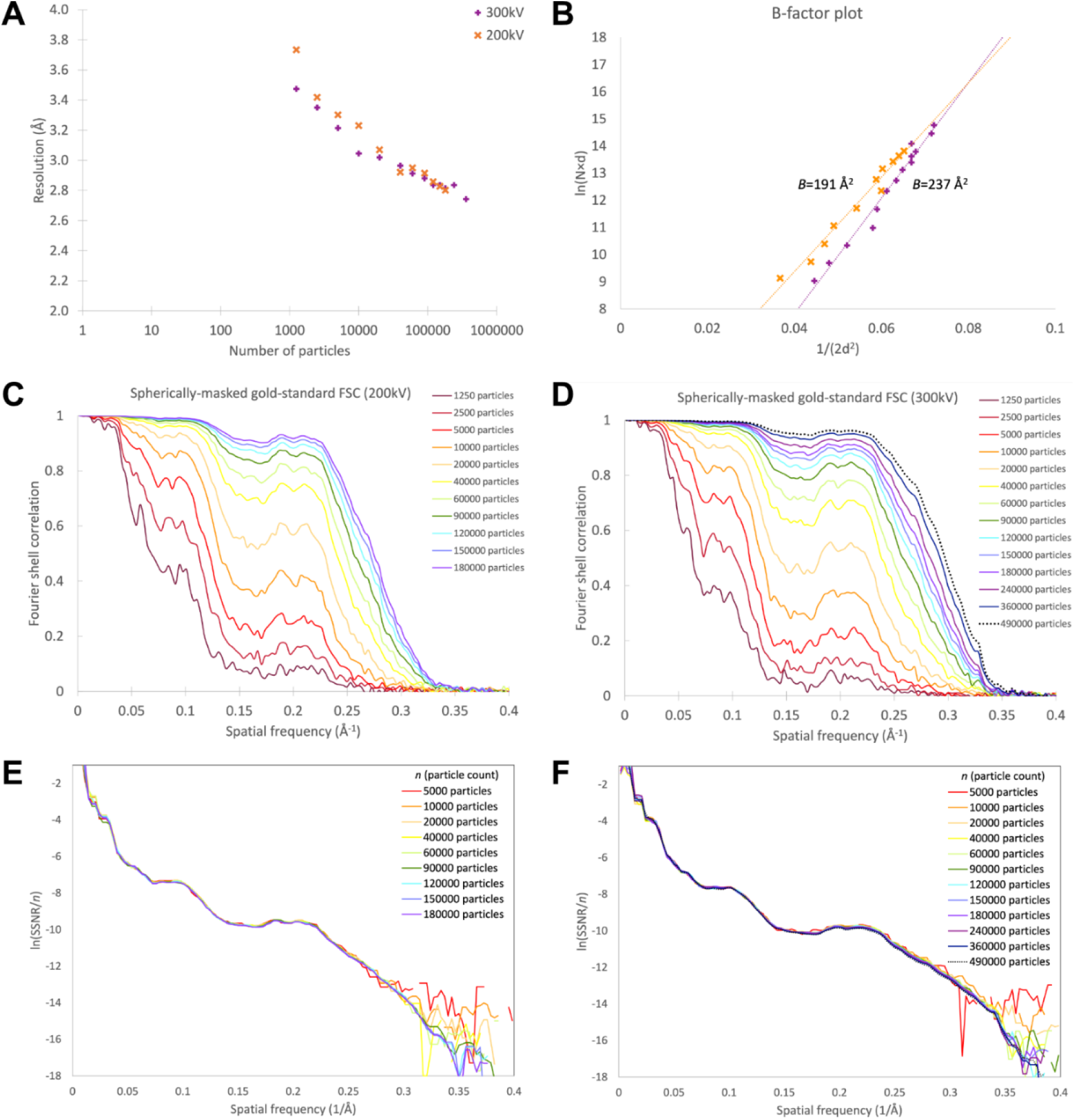
Effect of particle count on reconstructed information for *apo* ArnC. **A**) Resolution vs. number of particles for the dataset acquired with a 200 kV TEM (orange) and 300 kV TEM (purple). **B**) Transform of that data into a B-factor plot showing the natural log of the number of particles (N) multiplied by the resolution (d) plotted as a function of ½ d^-2^; colors as panel A. The slope of the B-factor plot represents the resolution-dependent signal dampening and is analogous to the crystallographic temperature factor. **C, D**) Spherically-masked gold-standard Fourier shell correlation between even-odd half-maps reconstructed with halves of the data subsets from data acquired at 200 kV (C) and 300 kV (D); colored by the total number of particles in the subset (i.e. both halves). **E,** F) Transform of that data into a “universal plot” for 200 kV (E) and 300 kV (F) datasets by dividing the Fourier shell correlation from panels (C) and (D) by the particle count and log-transforming. The “universal plots” illustrate the information content per particle.

**Figure S7.**
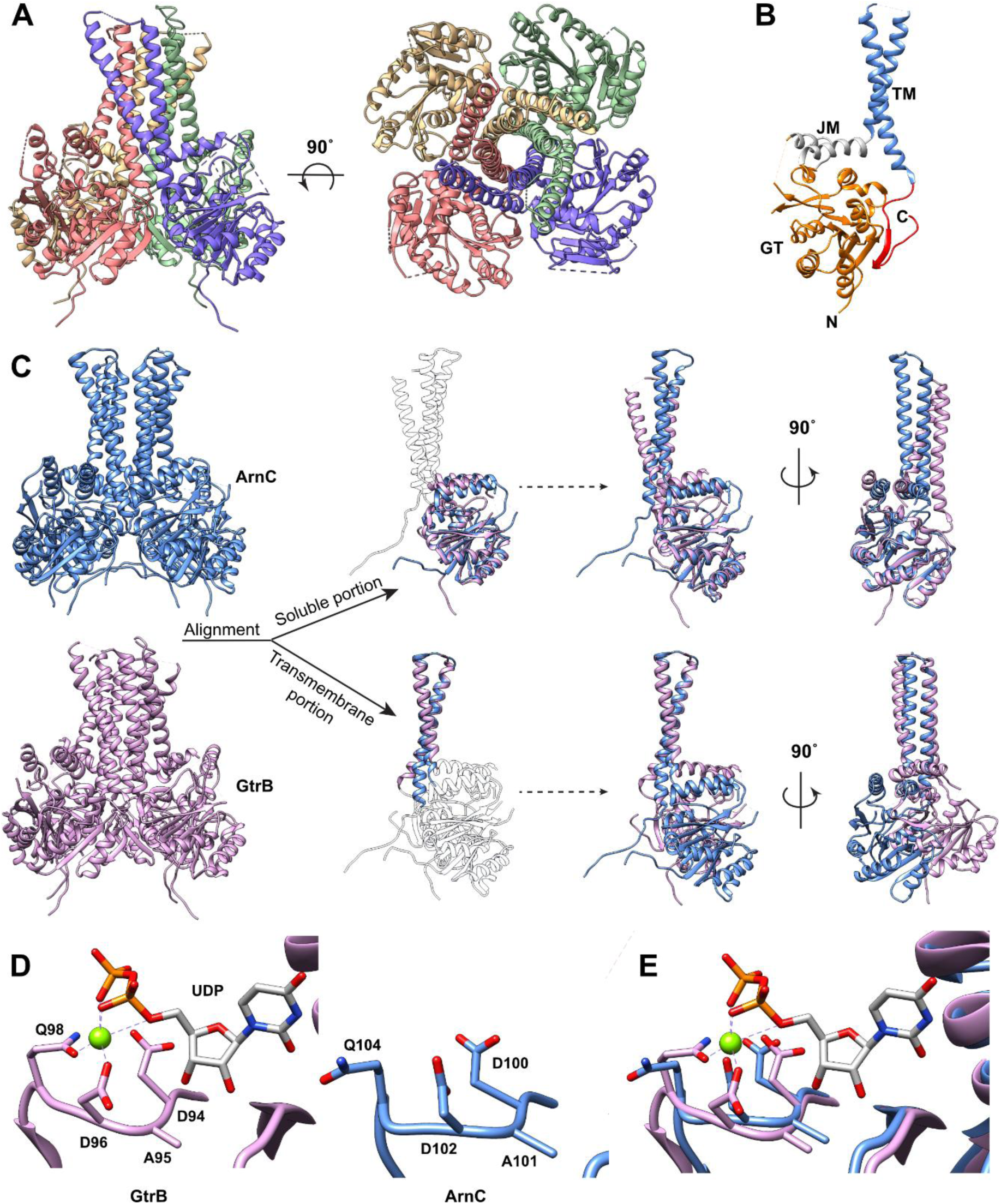
Comparison with the glycosyltransferase GtrB. **A**) Crystal structure of GtrB (PDB 5EKP) in ribbon representation with per protomer coloring. **B**) A single GtrB protomer is comprised of a GT-A fold glycosyltransferase domain (orange), two amphipathic juxtamembrane (JM) helices (gray), two transmembrane (TM) helices (blue) and a C-terminal β-hairpin (red). **C**) Comparison of ArnC (blue) and GtrB (pink), with a single protomer from each structure aligned via the GT-A soluble domain or the transmembrane domain. **D**) Comparison of the signature metal-coordinating DXD motif from GtrB (left) and *apo* ArnC (right). **E**) Superposition of the DXD motifs from GtrB and ArnC. Coloring as in (D).

**Figure S8.**
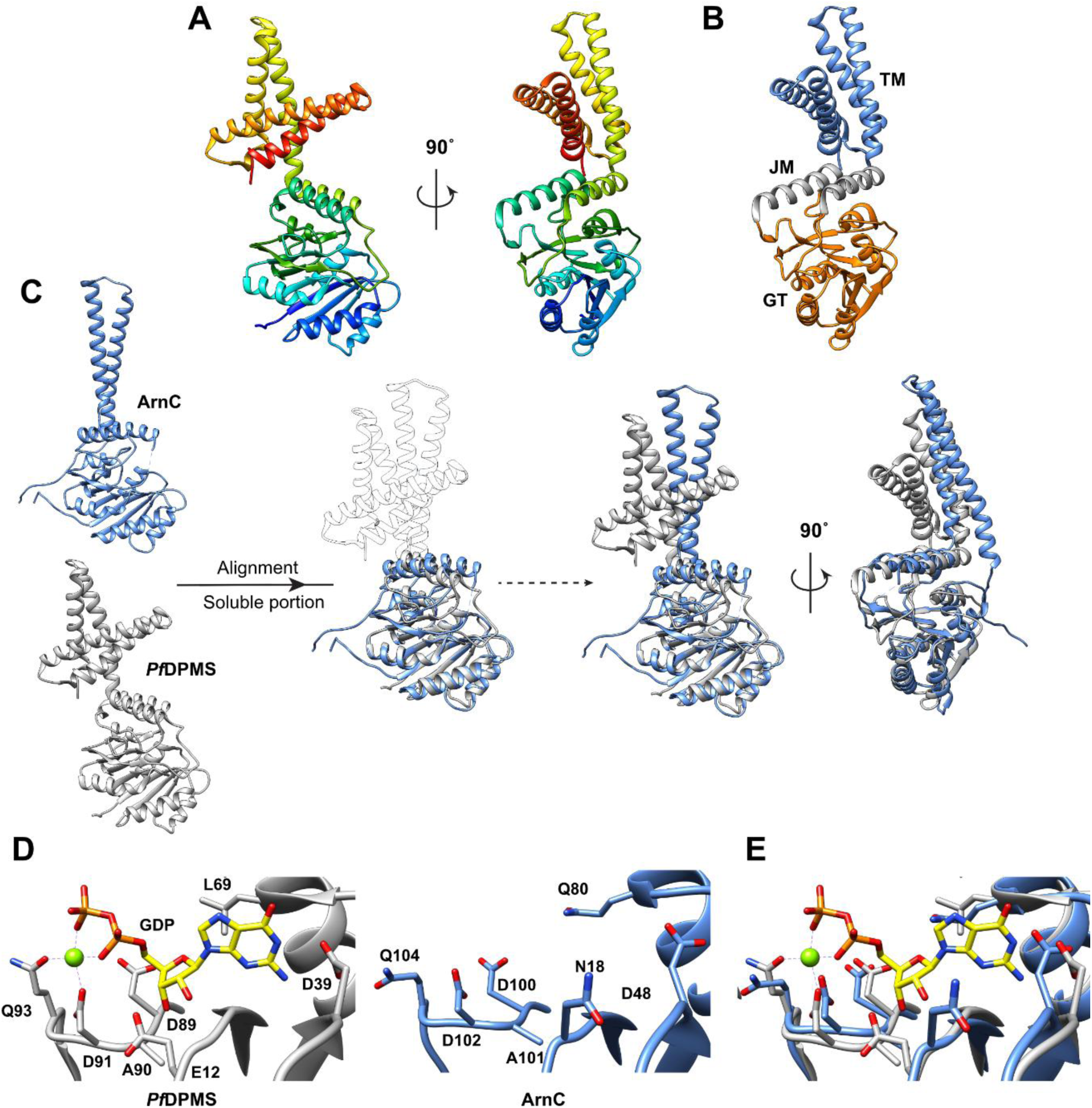
Comparison with *Pf*DPMS. **A**) Crystal structure of *Pyrococcus fiuriosus* DPMS (PDB 5MLZ) drawn in ribbon representation with rainbow coloring from N-terminus (blue) to C-terminus (red). Two orthogonal views are shown. **B**) DPMS is comprised of a GT-A fold GT domain (orange), two amphipathic juxtamembrane (JM) helices (gray), and four transmembrane (TM) helices (blue). **C**) Comparison of ArnC (blue) and DPMS (gray), with a single protomer of ArnC aligned with DPMS via the GT-A soluble domain. **D**) Comparison of the signature metal-coordinating DXD motif from DPMS (left) and *apo* ArnC (right). **E**) Superposition of the DXD motifs from DPMS and ArnC. Coloring as in (D).

**Figure S9.**
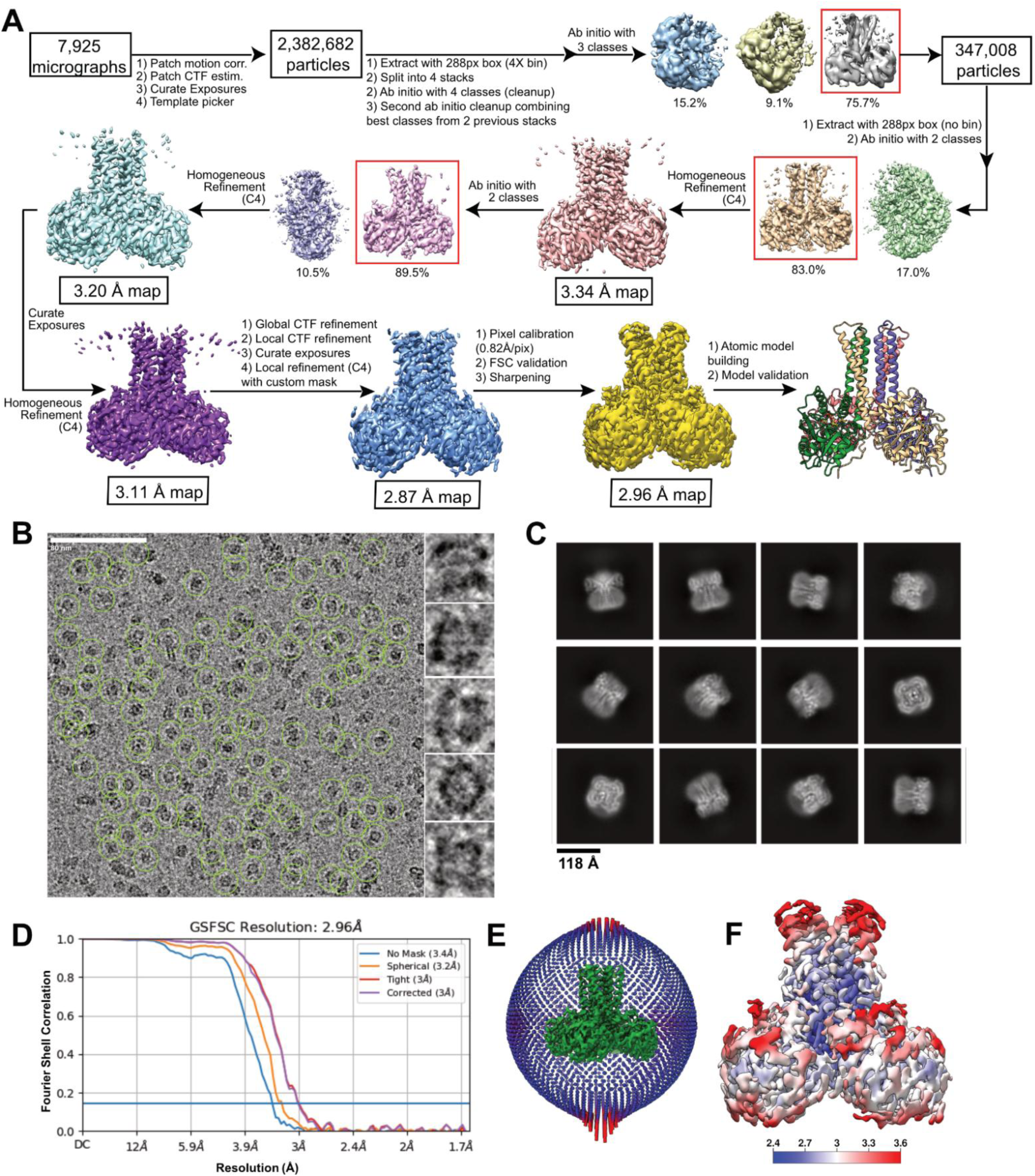
Cryo-EM analysis of UDP-bound ArnC*_Se_* collected on Talos Arctica. **A**) Data processing workflow used to determine the structure of nanodisc-reconstituted UDP-bound ArnC*_Se_* from the dataset collected on a Talos Arctica microscope. **B**) Representative electron micrograph of UDP-bound ArnC*_Se_* from Talos Arctica. Particles included in the final reconstruction are marked with green circles. Insets on the right show individual single particles from the micrograph. Scale bar, 80nm. **C**) Representative 2D class averages after reference-free 2D classification of the final particle stack in cryoSPARC. **D**) Fourier shell correlation (FSC) curve for UDP-bound nanodisc-reconstituted ArnC*_Se_* after the final local refinement in cryoSPARC. **E**) Euler angle distribution plot of all UDP-bound ArnC*_Se_* particles used in the final reconstruction. Final map shown in green. **F**) Local resolution map for the final UDP-bound ArnC*_Se_* reconstruction. Coloring shown from deep blue (2.4 Å) to red (≥3.6 Å).

**Figure S10.**
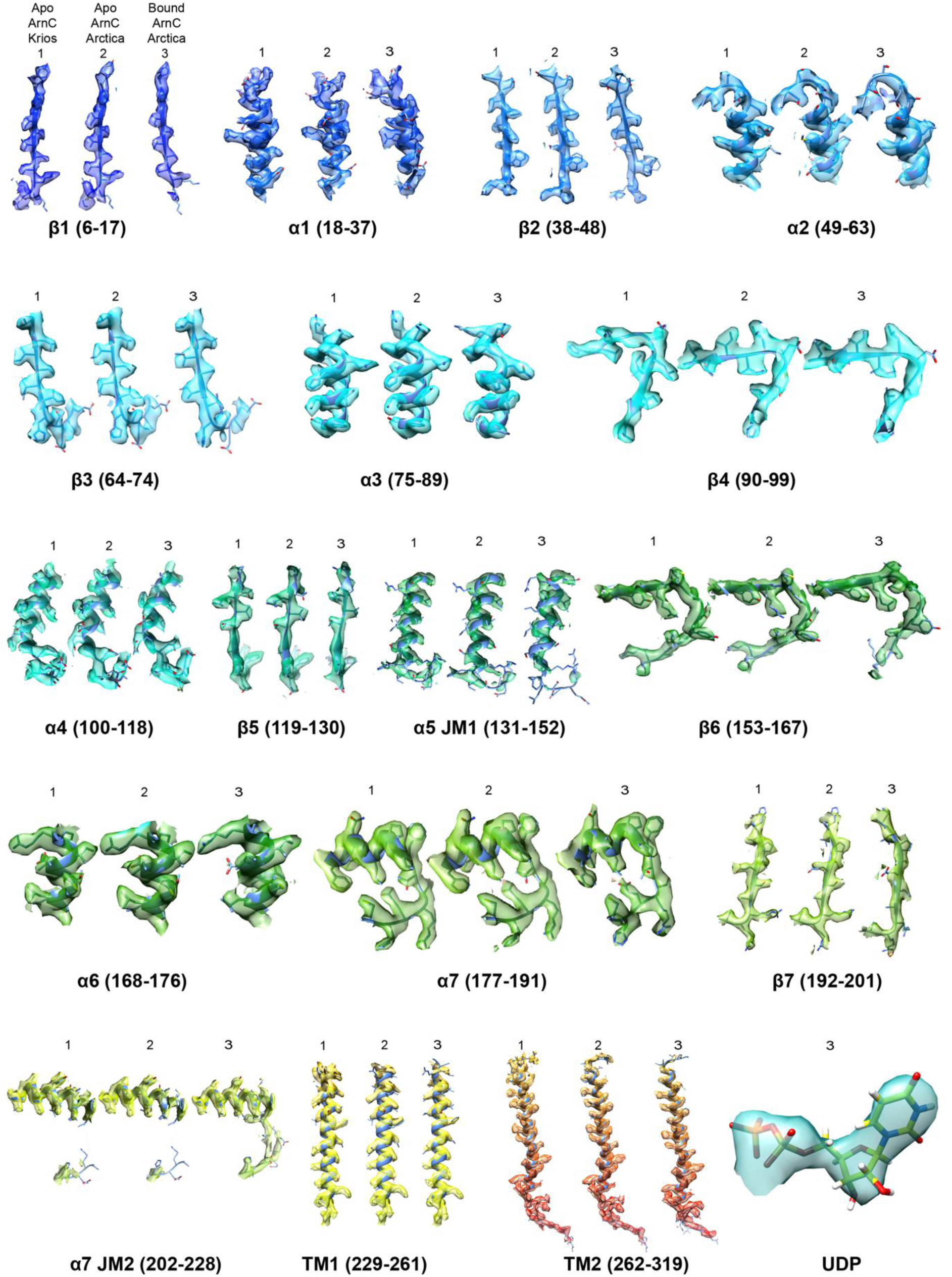
Cryo-EM densities of ArnC reconstructions. Cryo-EM densities in surface representation are superimposed on ArnC models for each of the three datasets as indicated: apo ArnC – Krios dataset (left), apo ArnC – Arctica dataset (middle), and UDP-bound ArnC – Arctica dataset (right). Coloring of densities is in rainbow from N-terminus (blue) to C-terminus (red). UDP (green) is shown as sticks.

**Figure S11.**
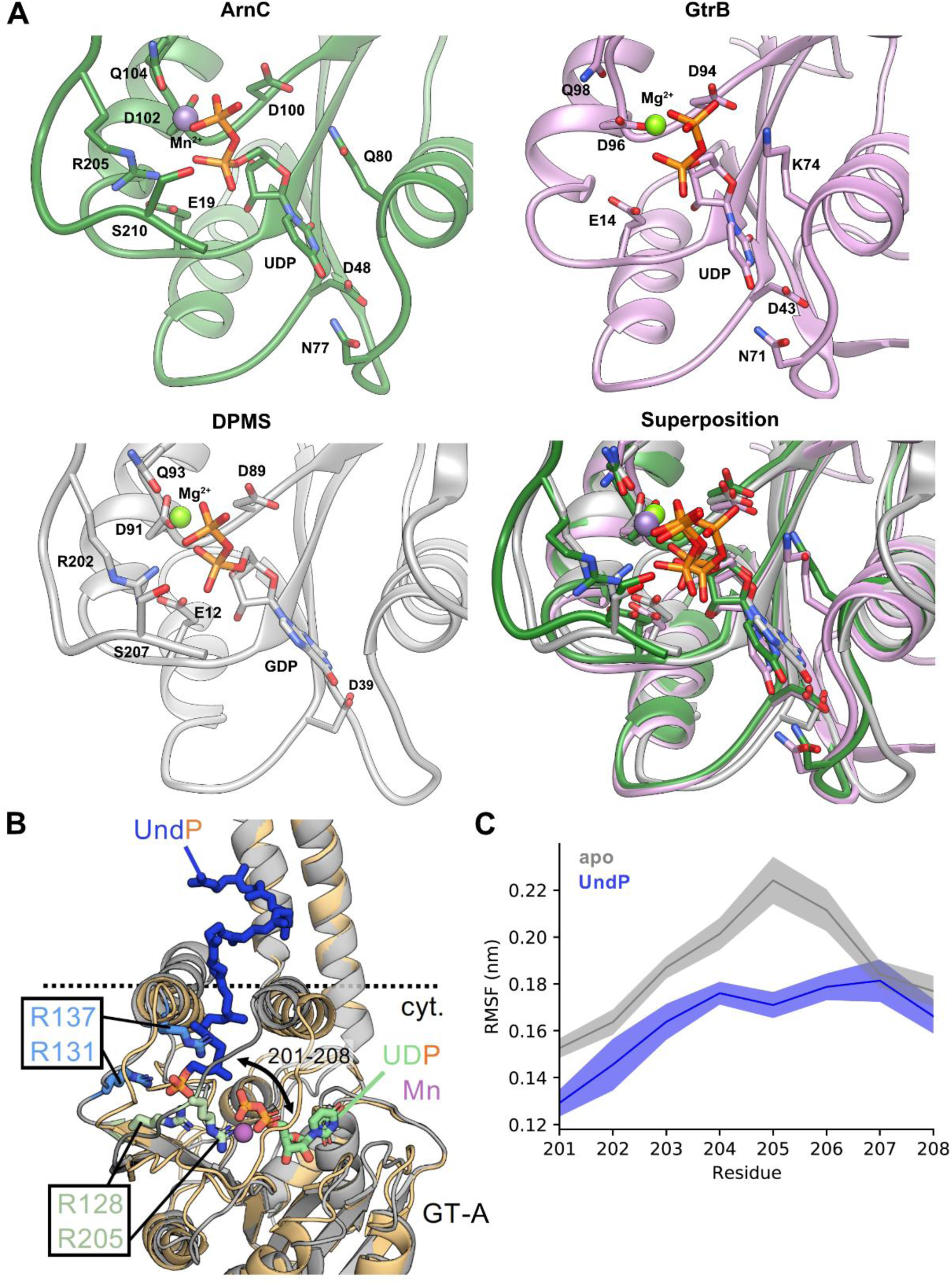
Substrate binding in ArnC. **A)** Comparison of nucleotide binding between the UDP-bound ArnC structure, GtrB (PDB 5EKP) and DPMS (PDB 5MLZ). Side chains for residues likely contributing to substrate binding (within 5Å of the nucleotide or the metal) are shown as sticks. Mn^2+^ is shown as a purple sphere and Mg^2+^ is shown as a green sphere. The superposition indicates the overlapping coordination of the nucleotide in all three enzymes. **B)** Snapshot of the final UndP binding pose within the ArnC GT-A domain from atomistic MD simulations (ArnC: gray, UndP: blue) overlayed with the ArnC-UDP structure (ArnC: light orange, UDP: green). UndP and UDP are shown as sticks and phosphate groups are colored orange. The Mn^2+^ ion is shown in purple. Arginine residues coordinating the UndP in simulations are shown in blue. Additional arginine residues facing the UDP site, and which may therefore be involved in coordination of incoming nucleotides, are shown in mint. UndP and UDP can both be coordinated within the GT-A domain without coordination clashes. The position of a flexible loop (residues 201-208) is indicated. **C)** Stabilization of the flexible loop by UndP. Mean root mean square fluctuation (RMSF) of the loop indicated in a across 3 x 100 ns atomistic simulations of ArnC with UndP bound to one GT-A domain (blue) compared to the three remaining apo GT-A domains (gray). Error bars represent standard error of the mean (SEM).

**Figure S12.**
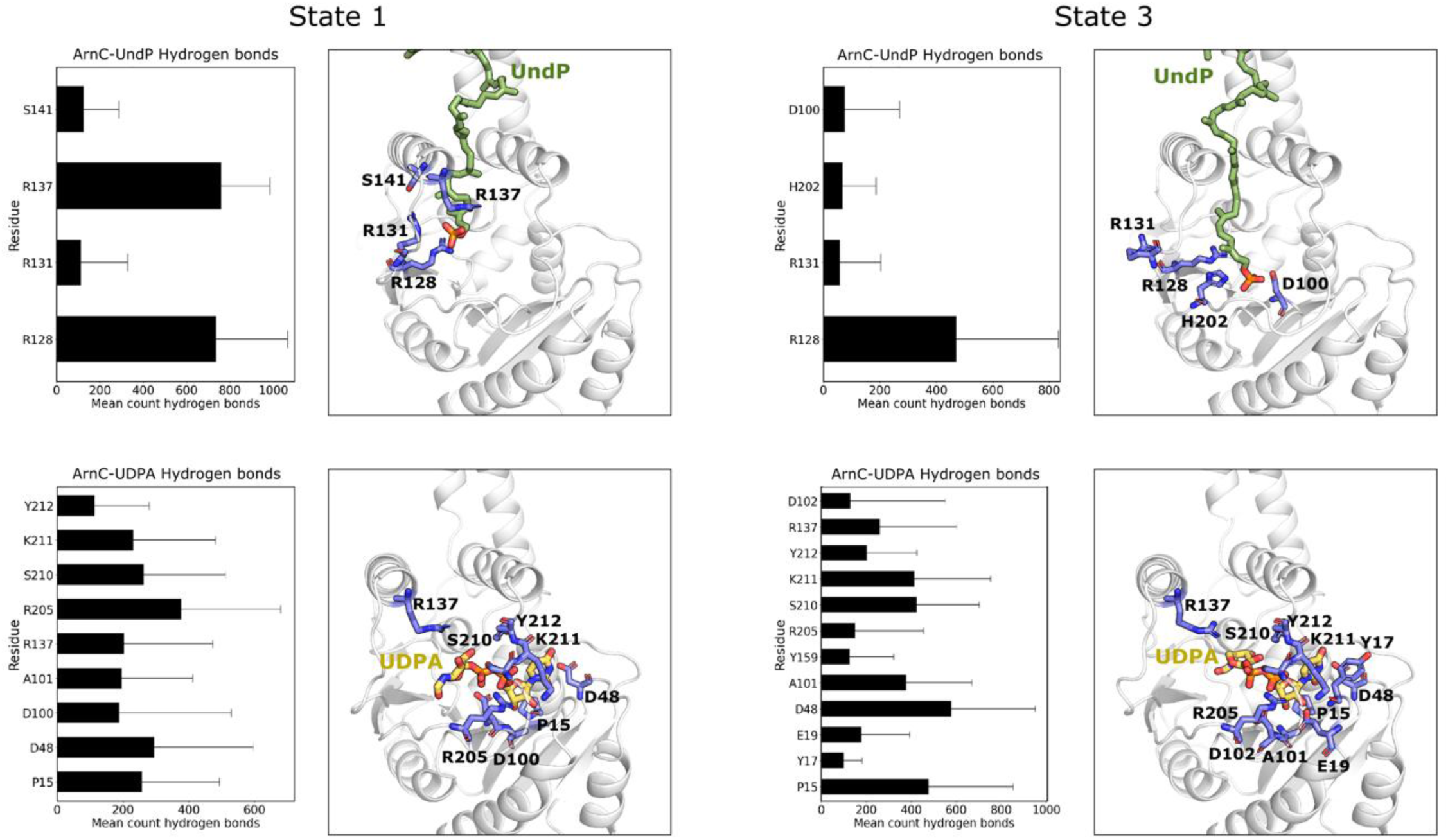
Hydrogen bonds between ArnC and its substrates. For each state, hydrogen bonds with undecaprenyl phosphate (UndP) (top) and UDP-L-Ara4FN (UDPA) (bottom) are shown. In state 1, the conserved arginines R128 and R137 coordinate the phosphate group of UndP, however, for state 3, only R128 contributes in the coordination of the lipid. Although the initial configuration for UDP-L-Ara4FN (UDPA) is different for state 1 and 3, its interaction with ArnC is similar for both states, due to the reorientation that the sugar group undergoes in state 1. Hydrogen bonds between the Ara4FN group and the conserved arginine R137 stabilize the sugar group for both states.

**Figure S13.**
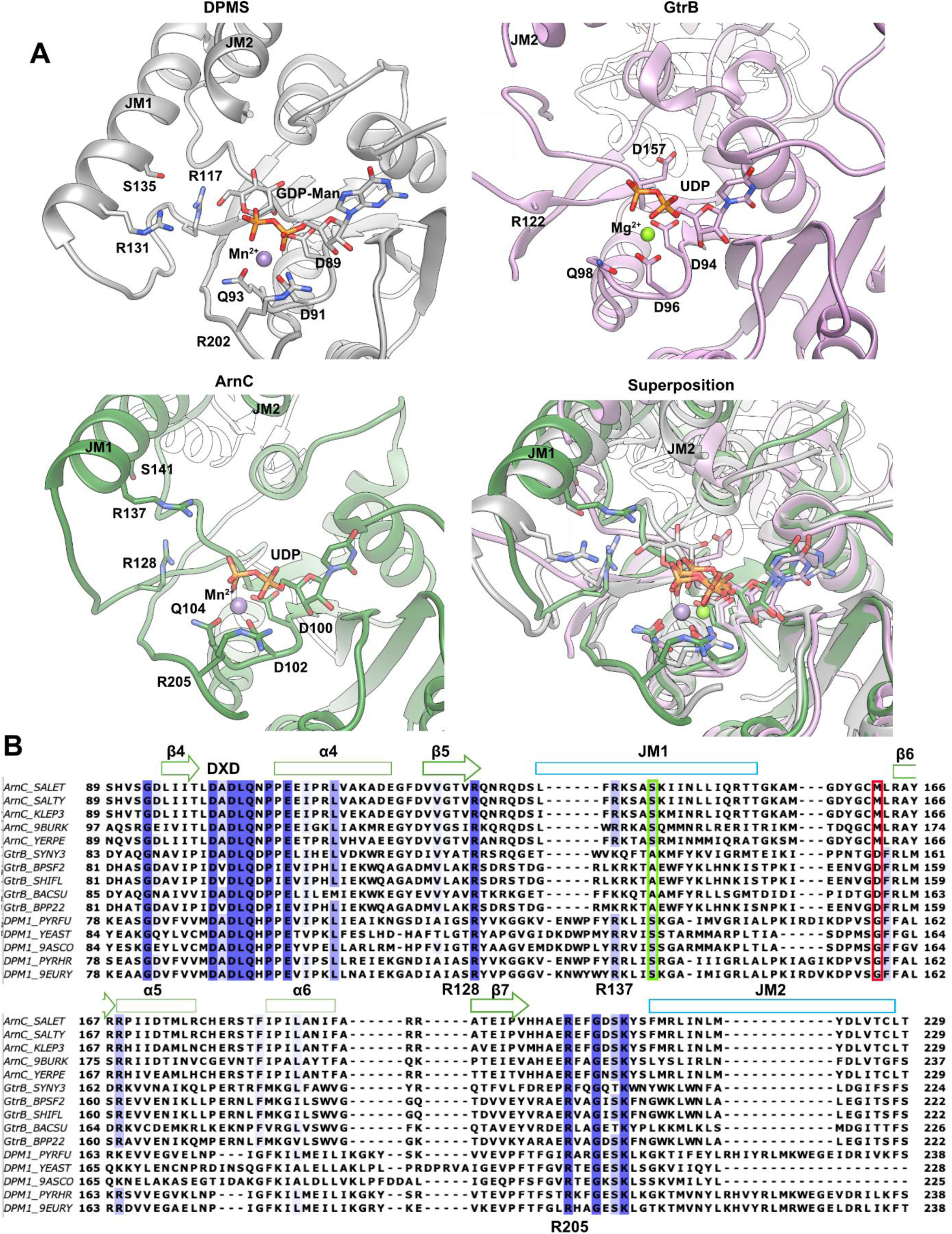
Catalytic residues in the ArnC, GtrB and DPMS families. **A**) Potential catalytic residues for DPMS (PDB 5MM0) (top left), and GtrB (PDP 5EKP) (top right), and their equivalent residues (based on position and conservation) in ArnC in the UDP-bound state (bottom left). A superposition of all three is shown in bottom right. Mn^2+^ shown as purple sphere, and Mg^2+^ shown as green sphere. The nucleotides (GDP-Man and UDP) are color matched to each structure with the phosphates shown in orange. **B**) Sequence alignment between β4 and JM2 of the GT-A domain from representative species of the ArnC, GtrB and DPMS families. Highly conserved residues are highlighted in blue and partially conserved residues are shown in lighter blue. *Pf*DPMS S141 and its corresponding residues in ArnC (Ser) and GtrB (Ala) families are highlighted by a green box. GtrB D157 and its corresponding residues in ArnC (Met) and DPMS (Gly) families are highlighted by a red box. The DXD motif and ArnC residues R128, R137 and R205 are marked on the alignment.

**Figure S14.**
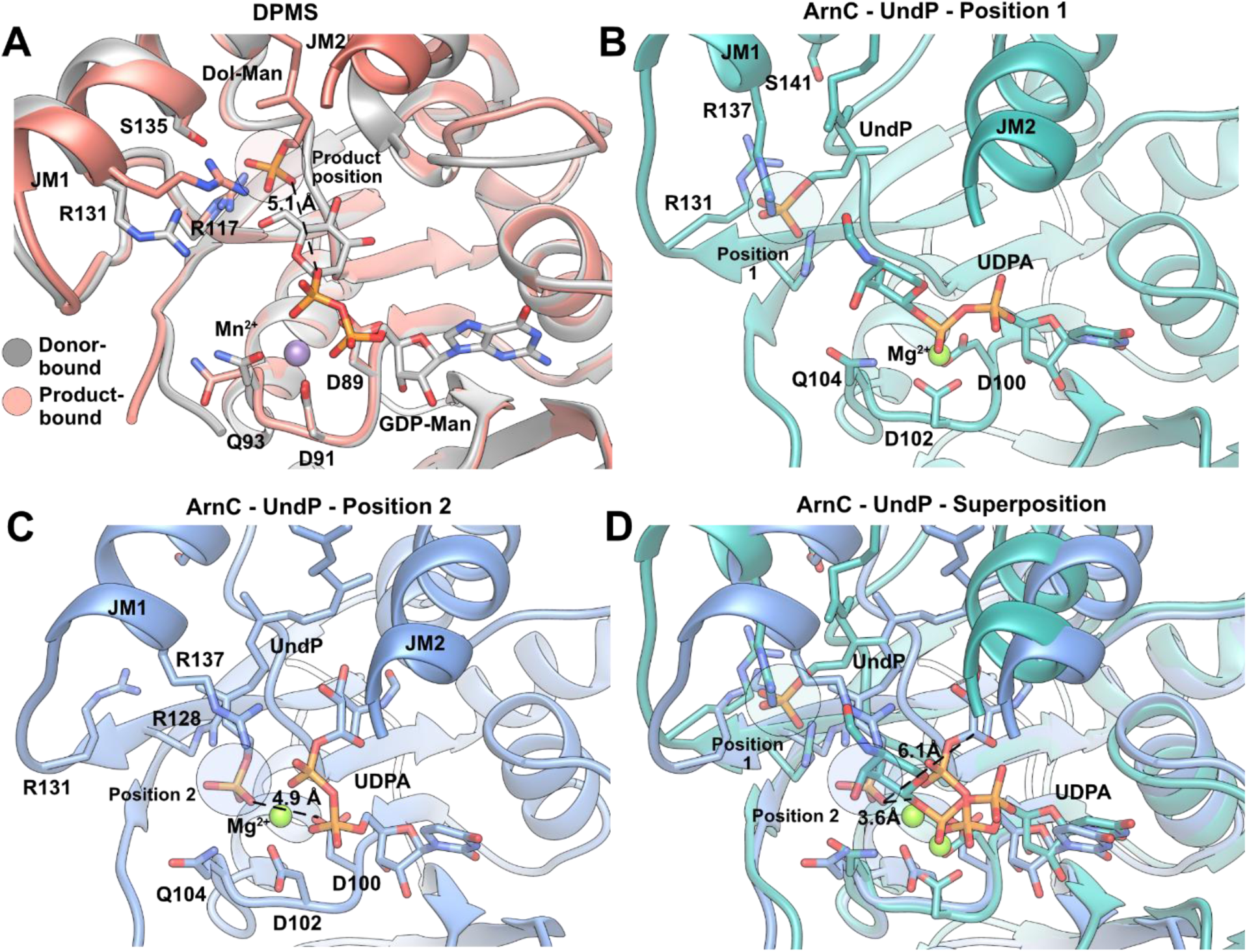
Coordination of the acceptor phosphate and catalysis in DPMS and ArnC. **A**) Superposition of the donor-bound *Pf*DPMS structure containing GDP-mannose (GDP-Man) (PDB 5MM0) and the product-bound *Pf*DPMS structure containing dolichol phosphate-mannose (Dol-Man) (PDB 5MM1). The mannose ring has been removed from the product-bound structure to showcase the position of the acceptor phosphate, which is highlighted with a red circle as the “Product position”. The distance between the nearest O atom of the acceptor phosphate and the C1 of the mannose ring in the donor-bound structure is 5.1Å. The Mn^2+^ of the donor-bound structure is shown as a purple sphere. **B**) Detail of the GT-A domain of ArnC from atomistic simulations corresponding to “State 1”, showing the acceptor phosphate in “Position 1” coordinated by R128, R131, and R137. In this position, the C1 atom of the donor L-Ara4FN sugar (UDPA) is 7.7Å away from the nearest O atom of the acceptor phosphate and is occluded by the rest of the sugar. **C**) Detail of the GT-A domain of ArnC from atomistic simulations corresponding to “State 3”, showing the acceptor phosphate deeper into the GT-A domain in “Position 2”, coordinated mainly by R128, Q104, D102. In this position, D100 is located 4.9Å away from the acceptor phosphate. **D**) Superposition of ArnC from (C) and (D). The superposition shows the two positions for coordination of UndP and highlights the distances between the nearest O atom of the acceptor phosphate in position 2 with the C1 atom of the L-Ara4FN sugar (3.6Å in “State 1”, and 6.1Å in “State 3). The Mg^2+^ ion in panels (B)-(C) is shown as a green sphere.

## Notes

https://doi.org/10.2210/pdb8vxh/pdb

https://doi.org/10.2210/pdb9b77/pdb

https://doi.org/10.2210/pdb9asc/pdb

### Competing Interest Statement

The authors have declared no competing interest.

